# NF-κB transcriptionally enhances p53 accumulation dynamics hampering DNA repair

**DOI:** 10.64898/2026.02.24.707448

**Authors:** Emanuele Colombo, Sara Pozzi, Alessia Loffreda, Francesca Genova, Erika Aloi, Timothy Heinichen, Paola Falletta, Matteo Mazzocca, Tom Fillot, Daniela Gnani, Alessandra Agresti, Marco E. Bianchi, Samuel Zambrano, Davide Mazza

## Abstract

Cells integrate multiple, often concurrent signals through intertwined genetic circuits whose dynamics shape transcriptional programs and cell fate decisions. Among these, the tumor suppressor p53 and the inflammatory transcription factor NF-κB are central regulators of stress responses in normal and cancer cells, yet their dynamic crosstalk under co-activation remains poorly characterized. Here, we combine genetic approaches, live cell imaging, transcriptomic analysis and mathematical modeling to dissect their dynamic interplay. We find that co-activation of NF-κB by the inflammatory cytokines TNF-α and IL-1β significantly enhance p53 nuclear accumulation upon genotoxic stress or Nutlin3a, and this effect is absent in NF-κB-deficient cells. Mechanistically, we show that cytokines induce an NF-κB-mediated increase of *TP53* transcription, and mathematical modeling indicates that it is sufficient to account for the observed increased p53 accumulation. Functionally, NF-κB co-activation rewires p53-dependent transcriptional programs and impairs p53-mediated DNA repair following genotoxic stress, due to a shift of p53 dynamics from oscillatory to more sustained accumulation; p53 oscillatory dynamics and DNA repair remain largely unaltered in absence of NF-κB. Our results uncover an amplification of p53 response in presence of inflammatory cues that is transcriptionally mediated by NF-κB and that results, counterintuitively, in functional antagonism.

**SIGNIFICANCE STATEMENT:** p53 dynamics have been shown to correlate with the cellular responses of cancer cells to genotoxic insults such as those delivered by chemo- and radiotherapies. However, these dynamics have been mostly studied in settings that do not account for the pro-inflammatory cues that cancer cells might receive from the microenvironment. By addressing this gap, our study uncovers a previously unappreciated mechanism by which inflammatory signals induce an increased p53 accumulation, leading to reduced DNA repair capacity. Our results provide mechanistic understanding on the origin of p53–NF-κB antagonism and show how dynamically interconnected signalling pathways can produce counterintuitive functional outcomes. Translationally, the contribution of inflammation to the sensitivity of normal and cancer cells to DNA damage might be exploited in the management of chemo- and radiotherapies.

## INTRODUCTION

Cells control gene expression in response to stimuli through genetic circuits (1) that shape transcription factors (TFs) dynamics leading to the timely activation of adequate transcriptional programs (2, 3). Moreover, increasing evidence links specific and alternative modes of TF dynamics in response to external cues to specific biological outputs (4). Since cells are typically exposed to multiple stimuli that might lead to the activation of different TFs, it is plausible that the mutual influence (crosstalk) between genetic circuits has evolved to modulate their dynamics and lead to specific outcomes. For instance, it has been shown that the proper synchronization between Notch and Wnt oscillatory TF signaling dynamics is fundamental for somite generation (5). Likewise, the nuclear localization dynamics of the mechanotransducer YAP/TAZ controls the dynamics of Nanog and Oct4 pluripotency TFs, determining stem cell fate decisions (6). Further, the cell cycle and circadian rhythms act as coupled oscillators (7) that set the rhythm for a plethora of physiological processes (8). Hence, unveiling how genetic circuits dynamically influence each other can provide important clues on how cells and organisms respond to complex situations in physiological and pathological conditions.

Due to their central role as regulators of the cell’s response to stress, much effort has been devoted to understanding how the TFs p53 and NF-κB mutually influence each other (9–11). Different studies broadly point towards functional antagonism (10), by which the inactivation of one of them typically leads to the excessive activation of the other (12). This resonates with the prevailing view on the opposing functions of these TFs in cancer, with p53 acting as the guardian of the genome that activates cell cycle arrest and pro-apoptotic programs upon genotoxic stimuli (13,14), while NF-κB is a central player in inflammation (15) that drives pro-survival and pro-proliferation programs (16, 17). Yet other examples highlight that NF-κB and p53 can instead cooperate to establish specific inflammatory transcriptional responses (18–20). Thus, their crosstalk is likely more complex, and each factor may skew the cell fate towards either proliferation, cell cycle arrest or death in a context- and stimulus-specific manner. However, most of the studies focusing on the NF-κB/p53 crosstalk do not consider the dynamics of these TFs: p53 and the NF-κB family members - including the strong transcriptional activator p65 (that we will refer to NF-κB in what follows) - are controlled by negative feedback loops involving MDM2 (21) and the IκB inhibitors (22), respectively, which give rise to dynamic modulation of TF activation (including oscillations in their nuclear concentration), producing distinct dynamical patterns of target gene expression (23, 24).

Notably, NF-κB dynamics can influence life-death cell fate decisions (25), cell differentiation and stimulus-specific cellular responses (26–28). On the other hand, p53 dynamics have been shown to dictate the response to chemo- and radiotherapy (29, 30). Specifically, oscillatory p53 leads to efficient DNA repair as compared to a more sustained accumulation of nuclear p53 - obtained using the small molecule Nutlin3a - which leads to delayed DNA repair kinetics (31) and activation of terminal programs such as senescence or apoptosis (29, 30). However, this temporal dimension has largely been overlooked when trying to characterize the NF-κB/p53 crosstalk and its potential functional outcomes.

For these reasons, here we combined genetic approaches with live cell imaging and mathematical modelling to gain insights on the dynamic crosstalk between p53 and NF-κB. We found that in multiple cancer cell lines the simultaneous activation of NF-κB with TNF-α and p53 with Nutlin3a leads to higher nuclear levels of the latter compared to Nutlin3a alone. This change of p53 dynamics is confirmed through live cell imaging and also observed upon γ-irradiation, where p53 dynamics shifts from oscillatory to a more sustained nuclear accumulation when NF-κB is activated by the pro-inflammatory cytokines such as TNF-α and IL-1β, but remain unchanged when NF-κB (p65) is knocked out. Mechanistically, we found that the activation of NF-κB leads to increased transcription of the *TP53* gene; mathematical modeling shows that this increase can explain the observed increased p53 accumulation dynamics in presence of p53-activating stimuli. This shift in p53 dynamics rewires transcriptional programs with a specific transcriptional alteration of p53 target gene expression. Functionally, we find that p53-dependent DNA repair upon γ-irradiation is impaired when co-activated NF-κB perturbs p53 oscillatory dynamics; instead p53 dynamics and DNA repair remain largely unchanged if NF-κB is knocked out. In short, our work demonstrates that the p53 response to DNA damage in presence of inflammatory cytokines that activate NF-κB lead to an enhanced nuclear accumulation of p53 which in turn delays the DNA repair.

## RESULTS

### TNF-α enhances p53 activation in various cell lines

To investigate how NF-κB and p53 activation influence each other, we performed a time-resolved quantitative immunofluorescence in various cell lines carrying intact p53 and NF-κB pathways (breast cancer MCF-7, colorectal cancer HCT-116 and osteosarcoma U2OS). Cells were exposed to 10ng/ml TNF-α (a cytokine leading to NF-κB activation) and 10 μM Nutlin3a (a small molecule inducing p53 accumulation by inhibiting its interaction with the negative regulator MDM2), alone or combined (see **Methods**).

In these cell lines, NF-κB nuclear translocation occurred with similar dynamics when stimulated by TNF-α or TNF-α + Nutlin3a (**Figure 1A**), while treatment with Nutlin3a alone did not induce NF-κB activation, as expected.

**Figure 1.**
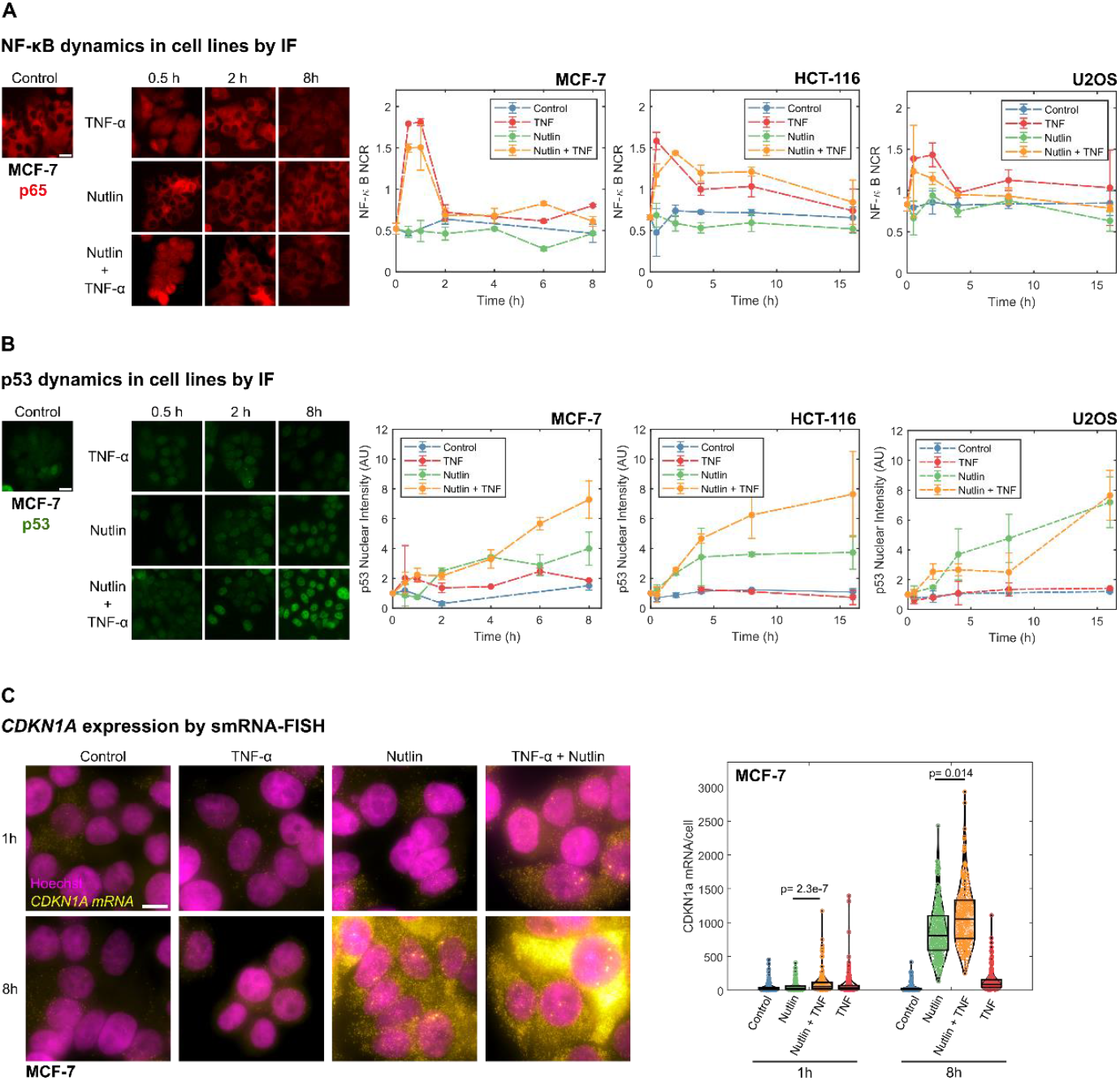
Co-activation of p53 and NF–κB leads to increased p53 accumulation. Representative images of immunofluorescence in MCF-7 cells against **A** p65 and **B** p53 at different timepoints and quantification of their nuclear to cytosolic ratio (NCR) and nuclear intensity, respectively, for MCF-7, HCT-116 and US2OS cells upon Nutlin3a, TNF-α and Nutlin3a+TNF-α treatments (scale bar 20 μm, errorbar: standard deviation, n = 2 biological replicates, with at least 150 cells per condition per replicate). **C** Representative images and quantification of smRNA-FISH against *CDKN1A* for MCF-7 cells 1h and 8 h post stimulation for Nutlin3a, TNF-α and Nutlin3a+TNF-α treatments (maximum projection of 3D stacks shown, scale bar 10 μm, n = 163, 188, 246, 233, 173, 190, 245, 180 cells for Control 1h, Nutlin3a 1h, Nutlin3a+TNF-α 1h, TNF-α 1h, Control 8h, Nutlin3a 8h, Nutlin3a + TNF-α 8h, TNF-α 8h respectively. Statistical test: Kruskal-Wallis, only relevant comparisons are shown for clarity).

Stimulation with TNF-α alone did not induce significant p53 activation in any of the cell lines tested. However, the dynamics of p53 upon Nutlin3a was strongly impacted by the simultaneous activation of NF-κB. Contrarily to our expectation of antagonistic activity of NF-κB and p53, co-stimulation with TNF-α and Nutlin3a resulted in a marked nuclear accumulation of p53 as compared to Nutlin3a treatment alone (an additional ∼40% increase at 16h, **Figure 1B**), both in MCF-7 and HCT-116, while no enhancement was observed in U2OS cells (**Figure 1B**). Western blot analysis in a panel of cell lines with WT p65 and p53 (**Figure S1**) further highlighted that co-treatment with Nutlin3a and TNF-α induced enhanced p53 accumulation in four of them (MCF-7, HCT-116 SKSH1, A549), three displayed similar p53 accumulation (U2OS, BJ and IMR-90), and in only one we observe a decreasing trend (NTera2).

To assess if the enhancement of p53 accumulation had functional consequences, we quantified the transcript levels of *CDKN1A* in MCF-7 cells at 1 and 8 hours post treatment by single-molecule FISH (smRNA-FISH) (32, 33) (**Figure 1C**). We observed a significant increase of *CDKN1A* mRNA upon co-stimulation with respect to Nutlin3a alone, but no direct upregulation of *CDKN1A* by stimulation with TNF-α, indicating that the observed increase in p53 levels can lead to an increased transcriptional output.

Taken together, our data show that co-stimulation with a p53 activator (Nutlin3a) and an NF-κB activator (TNF-α) has little effect on NF-κB dynamics but significantly enhances p53 nuclear accumulation leading to higher expression of the p53 target gene *CDKN1A*.

### Live cell imaging shows that p53 activation levels are enhanced by cytokines in an NF-κB dependent manner

To better characterize how TNF-α co-stimulation could enhance p53 activation, we resorted to live-cell microscopy. In the MCF-7 cell line, p53 displays rich dynamics that range from sustained activation upon Nutlin3a, to oscillations upon DNA damage by ionizing radiation (34). We therefore knocked-in GFP at the C-terminal of p53, obtaining the MCF-7-p53GFP cell line, as to preserve the endogenous regulation of the *TP53* gene (see **Methods** and **Figure 2A**). These cells were then further modified to either knock-out the NF-κB subunit p65 (NFκB^−^ cells) or to subsequently substitute it with a fluorescently tagged version (NFκB^+^ cells expressing tagged mScarlet-p65 only) (**Methods** and **Figure 2A-B**). We verified that the knockout for p65 in NFκB^−^ cells was successful (**Figure S2A**), that the p53-GFP knocked-in protein is equally upregulated by Nutlin3a in NFκB^+^ and NFκB^−^ cells **(Figure S2B)**, and that the tagged version of the TFs were capable of activating transcription of their target genes **(Figure S2C)**. Hence, we used live-cell confocal microscopy to quantify p53 and NF-κB dynamics in hundreds of cells (see **Methods, Figure 2A-B, Movie S1**) in response to TNF-α, Nutlin3a, or their combination.

**Figure 2.**
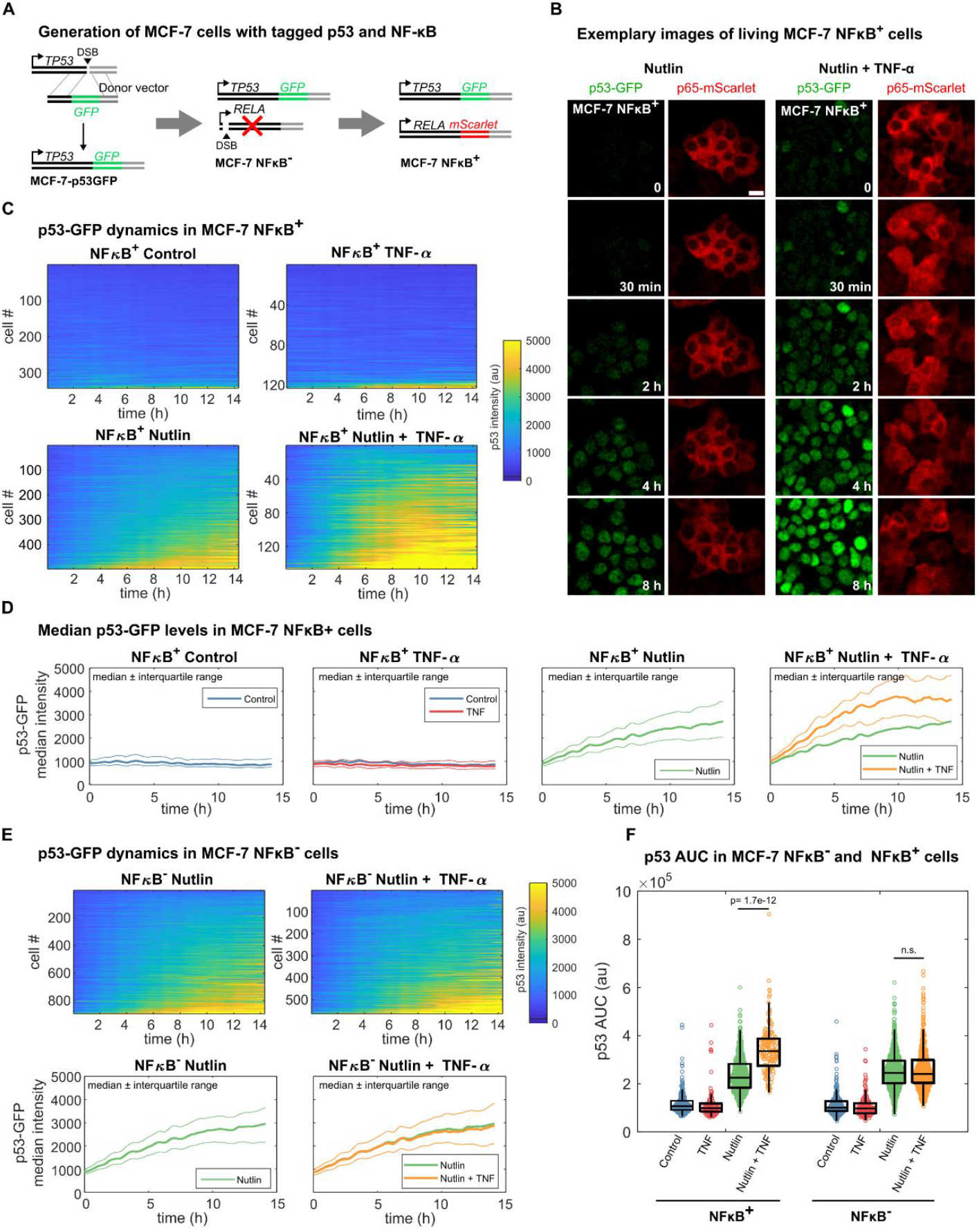
Co-treatment with TNF-α leads to enhanced p53 accumulation dynamics upon Nutlin3a in a NF-κB-dependent way. **A** Scheme of the generation of p53 knock-in, NFκB^−^ and NFκB^+^ MCF-7 cell lines. **B** Representative live cell imaging images of p53-GFP and p65-mScarlet in NFκB^+^ cells upon Nutlin3a and Nutlin3a+TNF-α treatments (scalebar 20 μm). **C** Colorplots of p53 activation dynamics for hundreds of NFκB^+^ single cells untreated, upon TNF-α, Nutlin3a, Nutlin3a+TNF-α and **D** representation of the median, first and third quartile (n = 342,121, 498, 146 cells respectively). **E** Colorplots of p53 activation dynamics in hundreds of NFκB^−^ single cells upon Nutlin3a and Nutlin3a+TNF-α treatments and representation of the median, first and third quartile (n = 354,149, 896, 564 cells for Control, TNF-α, Nutlin, Nutlin+TNF-α respectively). **F** Quantification and comparison of the AUC of NFκB+ and NFκB ^−^ cells upon Nutlin3a, TNF-α and Nutlin3a+TNF-α (Statistical test: Kruskal-Wallis, only relevant comparisons are shown for clarity).

Analysis of NF-κB dynamics in NFκB^+^ cells confirmed our previous observations: Nutlin3a alone did not induce NF-κB nuclear translocation (**Figure S2D**), TNF-α+Nutlin3a induced slightly lower NF-κB translocation compared to TNF-α alone, but both treatments induced similar NF-κB dynamics, with a first pulse of nuclear localization within the first hour post-stimulus (**Figure S2D**).

Live cell imaging of p53 dynamics were also well-aligned with our immunofluorescence results: in presence of functional NF-κB, Nutlin3a resulted in heterogeneous p53 accumulation dynamics, which was stronger upon co-stimulation with TNF-α (**Figure 2C-D**). Differently, treatment of cells knocked out for p65 (NFκB^−^, **Figure 2E**) with Nutlin3a alone or in combination with TNF-α resulted in comparable p53 dynamics, as quantified by the area under the curve (AUC, **Figure 2F**). This indicates that the observed enhancement of p53 accumulation upon co-treatment requires NF-κB. To further confirm that NF-κB activation enhances p53 accumulation, we co-stimulated our MCF-7 NFκB^+^ and NFκB^−^ cells with Nutlin3a and with another NF-κB-activating cytokine, IL-1ß: again, p53 dynamics were enhanced for NFκB^+^ cells and unperturbed for NFκB^−^ cells (**Figure S2E, F**).

Hence, live-cell imaging shows that there is a heterogeneous and yet enhanced accumulation of p53 upon coactivation of NF-κB and p53, in an NF-κB-dependent manner.

### NF-κB enhances p53 nuclear accumulation dynamics by upregulating *TP53* transcription

To gain insights on the molecular mechanisms of NF-κB mediated distortion of p53 dynamics, we first verified if the correlation between NF-κB activity and p53 response would hold at the single-cell level. Upon TNF-α or IL-1ß (**Figure S2G**) co-stimulation with Nutlin3a, cells displaying stronger NF-κB activation (see **Methods**) also displayed higher p53 response. Moreover, smRNA-FISH on the NF-κB and p53 targets *CDKN1A* and *NFKBIA* on single cells upon co-stimulation with Nutlin3a and TNF-α (**Figure S2H**) revealed a strong correlation between the expression of both targets, which further indicates that a strong NF-κB activation is accompanied by higher levels of p53. Taken together, our genetic models and single-cell imaging data support that the level of activation of NF-κB determines p53 response upon co-stimulation.

To characterize how NF-κB activation could affect p53 dynamics, we next assessed if NF-κB could alter the transcription of genes of the p53 regulatory circuit. The first natural candidate was the gene *TP53* itself, as it was previously reported as a potential NF-κB target (35, 36) and its promoter displays binding sites for NF-κB across vertebrates (see **Methods**). Indeed, by smRNA-FISH we could confirm that treatment with TNF-α or Nutlin3a + TNF-α resulted in significant increase of *TP53* mRNAs (up to 5-fold at 8 hours, **Figure 3A**); such upregulation is much less marked for NFκB^−^ cells, indicating that *TP53* transcription can be upregulated by p65 (**Figure S3A**).

**Figure 3.**
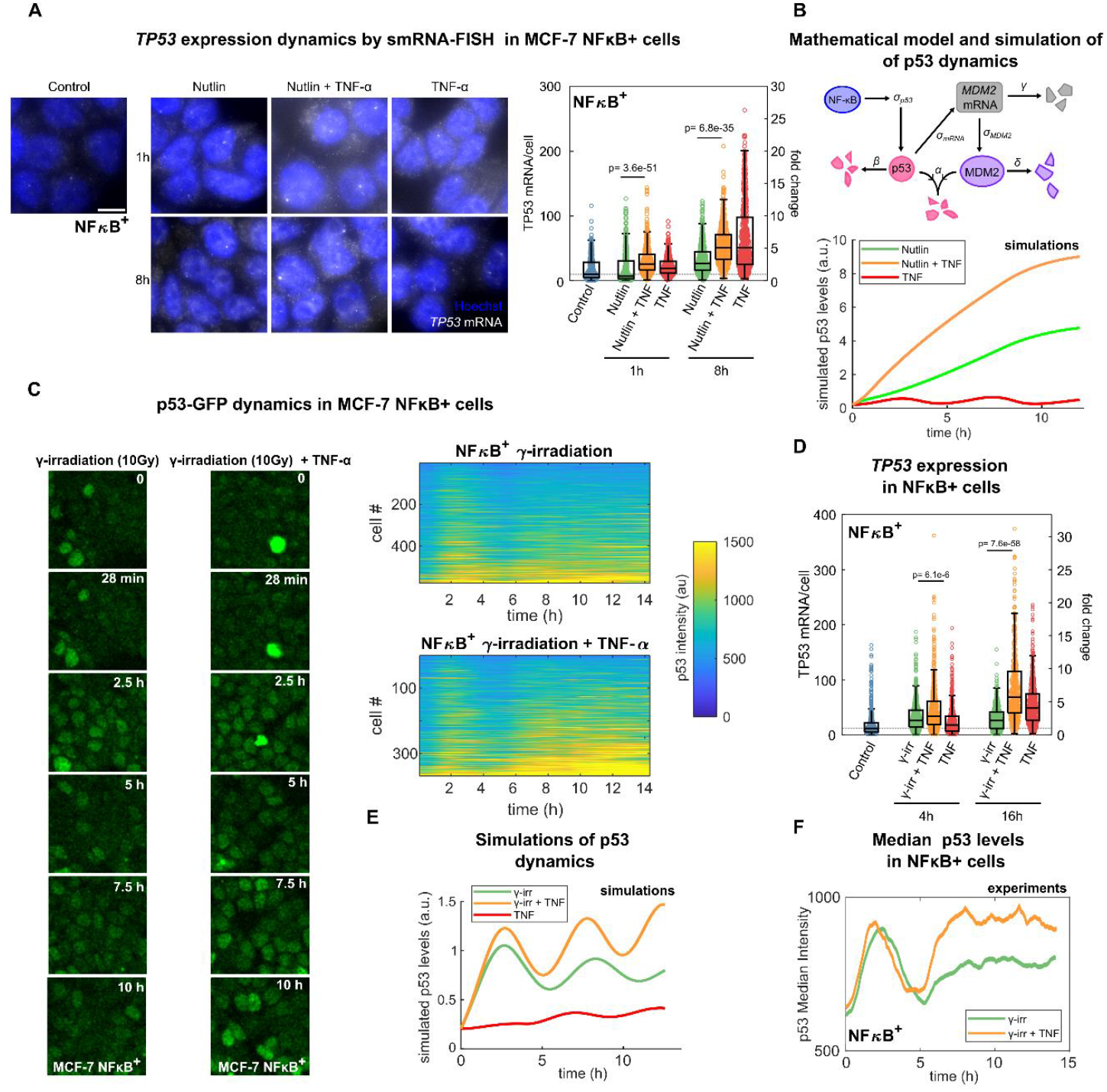
NF-κB upregulates *TP53* transcription and perturbs p53 oscillations in response to DNA damage. **A** Representative images of *TP53* expression through smRNA-FISH upon Nutlin3a and Nutlin3a+TNF-α treatments and quantification in NFκB^+^ cells (maximum projection of 3D stacks shown, scale bar 10 μm, n = 492, 607, 598, 639, 548, 594, 560 for Control, Nutlin3a 1h, Nutlin3a+TNF-α 1h, TNF-α 1h, Nutlin3a 8h, Nutlin3a+TNF-α 8h, TNF-α 8h, respectively. Statistical test: Kruskal-Wallis, only relevant comparisons shown for clarity). **B** Mathematical model of p53 regulation with key parameters, including the p53 synthesis rate, *σ*_*p*53_, modulated by NF-κB through increased TP3 expression (top). Simulations performed on p53 dynamics considering Nutlin3a alone, TNF-α alone and Nutlin3a + TNF-α applying the *TP53* modulations upon each treatment observed in A. **C** Representative images of p53 in NFκB^+^ cells upon γ-irradiation and γ-irradiation +TNF-α and colorplots representing the activation of hundreds of single cells (n = 580, 369 cells, respectively). **D** Quantification of *TP53* expression assayed through smRNA-FISH in NFκB^+^ cells untreated and upon TNF-α, γ-irradiation and γ-irradiation +TNF-α (n = 827, 698, 735, 632, 536, 443, 530 cells for control, γ-irradiation 4h, γ-irradiation+TNF-α 4h, TNF-α 4h, γ-irradiation 16h, γ-irradiation+TNF-α 16h, TNF-α 16h respectively). **E** Simulation of p53 oscillations for γ irradiation alone, TNF-α and γ+TNF-α considering time-modulated p53 synthesis following the *TP53* modulation values observed in D. **F** Median activation of p53 in NFκB^+^ cells upon γ-irradiation and γ-irradiation +TNF-α.

To gain quantitative insight on the effect of this increase in *TP53* in the context of the p53 regulatory circuit, we used a simple mathematical model (see **Methods**). The model builds on previous mathematical descriptions of the p53 response to DNA damage (24, 34, 35), that account for p53-mediated synthesis of MDM2 and MDM2-dependent degradation of p53 (**Figure 3B**). We first used the model to simulate the effect of Nutlin-3a: by reducing MDM2-mediated degradation of p53 we reproduced the dynamics of monotonic p53 accumulation observed in our cells (see **Figure 3B** and **Methods**). We next modified the model to explicitly incorporate a time-varying change in p53 synthesis rate *σ*_*p*53_, following our experimental observation of increased *TP53* expression (**Figure 3A-B** and **Methods**) upon Nutlin3a and TNF-α combined treatment. This single adaptation of the model in combination with the effect of Nutlin3a was enough to lead to an increase in the p53 nuclear accumulation dynamics compared to Nutlin3a alone (**Figure 3B**) in agreement with the one observed experimentally (**Figure 2C**). Of note, our simulations show that the enhancement in p53 accumulation requires both an increase in *TP53* synthesis and a reduction in MDM2-mediated p53 degradation (**Figure 3B**), consistent with our experiments showing that the TNF-α treatment alone does not lead to observable p53 accumulation (**Figure 2C**). Taken together, our data and modeling support that NF-κB activation affects p53 activation dynamics through an enhanced transcription of *TP53*.

### NF-κB activation perturbs p53 oscillations upon DNA damage

We next tested whether activation of NF-κB could also perturb p53 dynamics following a different stimulus, namely the induction of DNA damage by ionizing radiation (IR). Upon treatment of NFκB^+^ cells with 10Gy of γ-irradiation from a ^137^Cs source, p53 levels displayed an oscillatory behavior similar to what previously reported, with the first p53 peak observed at 2.5 hrs after stimulation (**Figure 3C and Movie S2**), and no discernible NF-κB activation (**Figure S3B**). Notably, smRNA-FISH revealed that co-stimulation with TNF-α and gamma rays did enhance *TP53* transcription (**Figure 3D**) up to 7-fold. We then went back to our mathematical model and evaluated what could be the effect of the resulting upregulation of the synthesis rate of p53, *σ*_*p*53_: we predicted that NF-κB co-activation would render IR-induced p53 oscillations faster with an increased accumulation of p53 (**Figure 3E**). Experimental results confirmed this prediction: TNF-α affected IR-induced p53 oscillations in NFκB^+^ cells, reducing their period and leading to a higher p53 nuclear accumulation (**Figure 3C, F**), as quantified by the timing of the second peak nuclear intensity reached during the 24 hrs time-course (**Figure S3C**) and by the AUC (**Figure S3D**), respectively. Also in this case, the observed perturbation of p53 dynamics by TNF-α depended on NF-κB (**Figure S3E**), since NFκB^−^ cells displayed less pronounced upregulation of *TP53* (**Figure S3F**) and less perturbed p53 oscillations upon co-stimulation with TNF-α (**Figure S3C**,**D**).

Overall, our results show that p53 oscillations upon γ-irradiation are significantly affected upon co-treatment with TNF-α and shifted towards a more sustained nuclear accumulation; this process is largely mediated by NF-κB-driven *TP53* transcription.

### Co-activation of NF-κB upon γ-irradiation reshapes transcriptional output and impairs DNA repair

We next verified how the shift in p53 dynamics translates into changes in the transcriptional response to stimuli.To this aim, we performed RNA-seq analysis in MCF-7 NFκB^+^ and NFκB^−^ cells in untreated conditions and at 4 and 16 hours after treatment with either γ-irradiation, TNF-α, or their combination. Principal component analysis (PCA) revealed that replicates corresponding to the same treatment clearly clustered together (**Figure 4A**),and highlighted more marked differences between treatments at the later timepoint. We therefore focused on the samples at 16h post-treatment for subsequent analysis. Differential expression analysis highlighted extensive transcriptional remodeling across all treatment conditions relative to untreated controls, consistent with broad activation of stress-responsive signaling programs (**Figure S4A**). As expected, gene set enrichment analysis comparing gamma-irradiated samples to untreated controls revealed significant upregulation of the p53 pathway in both NFκB^+^ and NFκB^−^ cells (**Figure S4B**), and enrichment of NF-κB-related pathways when comparing differentially expressed genes between NFκB^+^ and NFκB^−^ samples (**Figure S4C**). Both PCA (**Figure 4A**) and unsupervised clustering (**Figure S4D**) further showed that γ-irradiation and TNF-α induced predominantly opposing transcriptional responses, which resonates with the prevailing view of the antagonistic relationship between of p53 and NF-κB (10). Notably, this divergent gene-expression profile was largely preserved when comparing γ-irradiation alone with the combined γ-irradiation + TNF-α treatment. However, at the global scale p65 knock-out did not massively disrupt the transcriptional response to TNF-α, as NFκB^+^ and NFκB^−^ samples typically cluster next to each other (**Figure 4A and S4D**) This likely reflects the activation of additional transcriptional regulators downstream TNF-α, including non-canonical NF-κB dimers (e.g. encompassing c-Rel (39)) alongside with p38 and JNK kinases and pro-apoptotic pathways (40).

**Figure 4.**
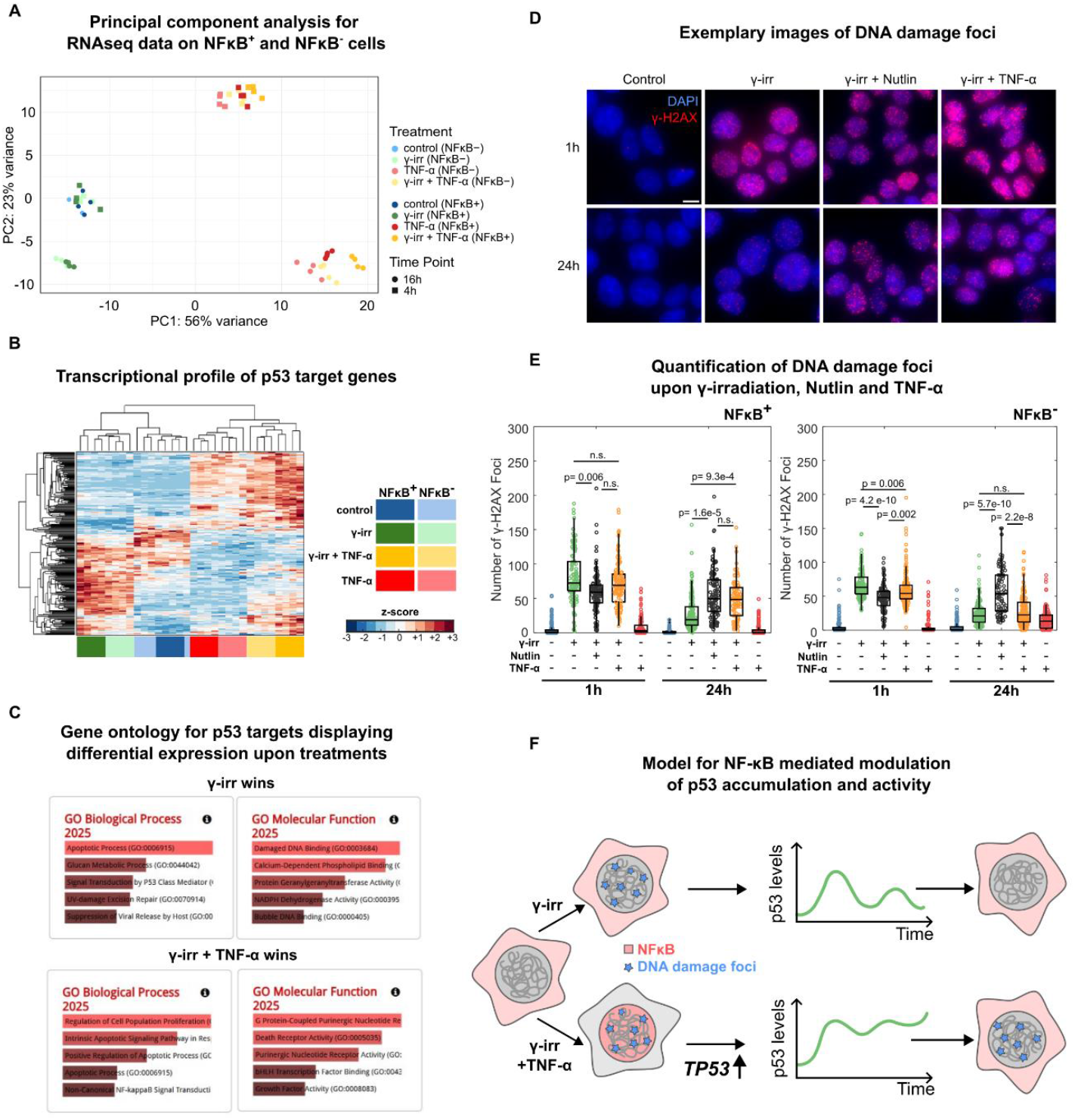
NF-κB activation upon γ-irradiation rewires the transcriptional response and hampers DNA repair. **A** Principal component analysis (PCA) of the RNAseq data of NFκB^+^ and NFκB^−^ cells upon no treatment or 4h and 16h treatment with TNF-α, γ-irradiation and γ-irradiation +TNF-α. **B** Hierarchical clustering of p53 targets for NFκB^+^ and NFκB^−^ cells under no treatment, or 16 hrs treatment with TNF-α, γ-irradiation and γ-irradiation +TNF-α. Z-score represented for genes differentially expressed in at least one condition (p<0.05 upon Bonferroni-Holmes correction). **C** Pathways with enriched gene ontologies in each gene set identified for “Biological Process” and “Molecular Function”. **D** Representative images of gamma-H2AX in NFκB^+^ cells after 1h and 24 h upon γ-irradiation, γ-irradiation +TNF-α, γ-irradiation and γ-irradiation+Nutlin3a (maximum projection of 3D stacks shown, scale bar 10 μm). **E** Quantification of the number of gamma-H2AX foci for untreated cells and upon γ-irradiation 1h, γ-irradiation+Nutlin3a 1h γ-irradiation+TNF-α 1h, control 24h, γ-irradiation 24h, γ-irradiation+Nutlin3a 24h, γ-irradiation+TNF-α 24h for NFκB^+^ cells (n = 245, 120, 129, 118, 275, 165, 108, 93 cells, respectively, statistical test: Kruskal-Wallis) and NFκB^−^ cells (n = 156, 168, 141, 188, 179, 157, 97, 158 respectively, statistical test: Kruskal-Wallis, only relevant comparisons shown for clarity). **F** Working model on the effect of combined NF-κB and p53 activation: NF-κB induces *TP53* expression, perturbing p53 oscillatory dynamics and hampering DNA repair.

Considering the observed shift in p53 dynamics following γ-irradiation, we next investigated how this specifically affected p53-dependent transcription. We analyzed the expression of a set of consensus p53 target genes (41) and clustered them according to their responses to the different treatments. Combined γ-irradiation and TNF-α treatment produce the most distinct pattern of p53 target gene expression pattern compared to γ-irradiation alone, an effect that was partially reduced by p65 knock-out (**Figure 4B**). To further characterize the functional consequences of this modulation we compared the gene expression changes induced by γ-irradiation alone with those induced by irradiation + TNF-α treatment. We identified a subset of genes more strongly induced by γ-irradiation alone (“γ wins” **Figure S4E**), which were enriched in biological terms related to a variety of p53 dependent mechanisms including DNA damage response and repair. In contrast, genes more strongly induced by the combined treatment (“γ + TNF wins”) were more skewed towards apoptotic processes (**Figure 4C**). Changes in expression of these genes upon-cotreatment were slightly more pronounced in NFκB^+^ cells than in NFκB^−^ (**Figure S4F**). We therefore hypothesized that the co-treatment with TNF-α can indeed change the efficiency of DNA repair upon γ-irradiation. This would fit both with the transcriptional effect of the co-treatment described above and recent theoretical and experimental work which highlights that sustained p53 dynamics might result in less efficient DNA repair than p53 oscillations (31).

We then monitored by immunofluorescence gamma-H2AX (**Figure 4D**) a marker that accumulates at DNA damage foci, following different stimuli: γ-irradiation (that results in oscillating p53 levels), γ-irradiation + TNF-α (that in presence of p65 perturbs the oscillatory dynamics increasing its nuclear accumulation) and γ-irradiation + Nutlin3a (that induces a monotonic increase of p53 independently of the NF-κB pathway, **Figure S4G**, as reported also by others (31, 34)). The number of gamma-H2AX foci was analyzed at 1hr to monitor the extent of DNA damage and at 24hr to assess DNA repair efficiency (see **Methods**).

In NFκB^+^ cells (**Figure 4E**), γ-irradiation induced similar DNA damage levels in all conditions (as probed 1hr post-irradiation). However, 24h post-irradiation, DNA damage is reduced to a greater extent for cells exposed to γ-irradiation alone (∼75% reduction) than to both the γ-irradiation+Nutlin3a (∼15% reduction) and the γ-irradiation+TNF-α combinations (∼28% reduction); this is in line with previous work showing that perturbation of p53 oscillatory dynamics leads to less efficient DNA damage repair (31) (**Figure 4E**). In contrast, in NFκB^−^ cells (**Figure 4E**), where co-treatment with cytokines does not affect the oscillatory p53 dynamics (**Figure S3E.F**), DNA damage is efficiently repaired in both cells treated with γ-irradiation or γ-irradiation + TNF-α (**Figure 4D**). Of note, also for these cells, the perturbed dynamics upon γ-irradiation + Nutlin3a leads to a much worse repair, further highlighting the role of oscillatory p53 dynamics in this process.

Finally, analogous results were obtained when stimulating NFκB^+^ and NFκB^−^ with IL-1ß: upon-costimulation with this cytokine and γ-irradiation, NFκB^−^ cells repaired DNA damage more efficiently than NFκB^+^ cells **(Figure S4H**).

Our results hence indicate that NF-κB activation upon γ-irradiation induces a rewiring of the transcriptional programs and an enhancement of p53 dynamics response that hampers DNA repair.

## DISCUSSION

Our work reveals that p53 dynamics are actively reshaped by inflammatory cues through NF-κB– dependent transcriptional control of *TP53*. NF-κB enhances *TP53* transcription which results in a shift of p53 nuclear dynamics from oscillatory to a more sustained accumulation, which in turn decreases the ability of cells to repair DNA damage. This crosstalk establishes a paradoxical relationship in which NF-κB cooperates with p53 at the molecular level while antagonizing its genome-protective function (**Figure 4F**).

### Inflammatory stimuli enhance p53 accumulation upon DNA damage, impairing DNA repair

It is now well established that cell-fate decisions following DNA damage are in part instructed by the dynamics of p53 nuclear accumulation (29, 34, 42). For example, pharmacological inhibition of MDM2 negative feedback with Nutlin3a converts p53 dynamics from oscillatory to sustained nuclear accumulation in irradiated cancer cell lines, delaying DNA repair (31) and biasing cells towards terminal outcomes such as senescence and apoptosis (34). However, exposure of cells to Nutlin3a is unphysiological, and it remained largely unexplored whether physiological inputs can similarly reshape p53 dynamics and thereby perturb the DNA damage response. Here, we have investigated this possibility by focusing on pro-inflammatory cytokines that activate the NF-κB pathway.

We show that co-stimulation with TNF-α or IL-1β markedly enhances p53 nuclear accumulation relative to treatment by Nutlin3a or γ-irradiation alone. This amplification requires NF-κB activity, as genetic ablation of the p65 subunit nearly abolishes cytokine-induced boosting of p53 levels. Notably, this regulatory interaction is largely unidirectional: NF-κB activation increases p53 accumulation, whereas NF-κB nuclear translocation dynamics are mostly insensitive to p53 activation. Functionally, the cytokine-driven shift towards increased p53 nuclear accumulation in NF-κB–competent cells impairs the repair of γ-irradiation–induced DNA damage, an effect that is again abolished by p65 knock-out.

While reinforcing the concept that p53 dynamics encode cell-fate information, our work reveals that such dynamics can be reprogrammed by crosstalk with NF-κB driven inflammatory signaling pathways.

### NF-κB and p53: molecular cooperation leading to functional antagonism

In cancer biology, NF-κB and p53 are traditionally viewed as antagonistic transcription factors that lead to opposing functional outcomes (10, 12). Consistently with this prevailing view, our transcriptomic analyses show that γ-irradiation and TNF-α elicit largely opposing genome-wide responses. Yet, beyond this global antagonism, we uncover a direct regulatory link whereby NF-κB promotes *TP53* gene expression, resulting in enhanced and more sustained p53 nuclear accumulation. However, this cooperative regulation can be reconciled with the notion of functional antagonism, since enhanced accumulation of p53 impairs DNA repair, thereby uncoupling p53 abundance from its canonical protective role in genome maintenance. Thus, while at the molecular level NF-κB activation enhances p53 nuclear accumulation, it paradoxically antagonizes p53 function at the level of DNA damage resolution. The ambivalent crosstalk observed here between NF-κB and p53 might contribute to explaining how both pathways can elicit pro-survival or pro-apoptotic responses in a context- and stimulus-dependent manner (4, 19, 20, 43, 44).

Though these results reinforce the recent findings highlighting that a shift of p53 oscillatory dynamics towards a monotonous increase compromises DNA repair (31), the precise mechanisms involved remain to be fully elucidated. Recent experimental and theoretical studies suggest that p53 may directly facilitate the recruitment of DNA damage response factors in *cis* at sites of DNA lesions (45), and that redistribution of p53 across multiple genomic binding sites may be sensitive to its nuclear accumulation dynamics. In this scenario, p53 dynamics may influence its spatial distribution through mechanisms such as phase separation and Ostwald ripening (31), providing a potential physical basis by which excessive or sustained p53 accumulation disrupts efficient coordination of the DNA damage response.

### NF-κB induced upregulation of *TP53* as a potential mechanism coupling inflammation to genome surveillance

We identify NF-κB as an upstream regulator of *TP53* expression. This is consistent with the reported ability of NF-κB to bind the *TP53* promoter (35, 36) and is functionally supported by our observation that cytokine-induced *TP53* transcriptional upregulation is largely abolished upon genetic ablation of the NF-κB p65 subunit. Mathematical modelling shows that NF-κB-mediated enhanced transcription of *TP53* can account quantitatively for the notable increase in p53 nuclear accumulation observed in our system upon co-stimulation with Nutlin3a or γ-irradiation and inflammatory cytokines. Of note, cytokines alone lead to NF-κB activation and increased *TP53* expression, but in absence of concomitant p53-activating stimulus this does not lead to observable changes in p53 levels; this is again recapitulated by our mathematical model of the p53 regulatory network. The substantial change in p53 accumulation dynamics hereby described was not observed in a recent study where cells were stimulated with γ-irradiation and TNF-α and this seemed to affect p53 oscillatory dynamics less dramatically (46); their use of a fluorescently tagged p53 under the control of an exogenous promoter or the cell-type specificity that we observe here could explain this discrepancy.

Notably, we do find that NF-κB binding sites within the *TP53* promoter are conserved across vertebrates (see **Methods**), pointing to evolutionary pressure to couple and coordinate inflammatory signaling with p53 upregulation. Indeed, in macrophages a precise cooperative transcriptional program between p53 and NF-κB has been identified (20) by which p53 contributes to amplify cytokines and chemokines transcription, further pointing to a more subtle and cell-specific regulation between these two TFs. We also note that cytotoxic T cells kill target cells by delivery of granzymes, which induces apoptosis and DNA breaks (47); our findings raise the possibility that T-cell attack efficacy might be facilitated by pro-inflammatory cues, as cytokine-driven NF-κB activation would lower the p53-dependent ability of target cells to repair such breaks. Our findings potentially have even broader implications: cancer cells are more prone to accumulate DNA damage, and an inflammatory environment could further hamper their ability to mount a proper response, pushing them toward apoptosis or senescence. Under this light, the use of NF-κB-inhibiting drugs like dexamethasone (48) could decrease therapeutic efficacy if applied concomitantly to radiotherapy (49) and the same might be true for chemotherapy.

In summary, our work reveals that p53 dynamics are actively reshaped by inflammatory cues through NF-κB–dependent transcriptional control of *TP53* and that this crosstalk impairs DNA repair. These findings suggest that inflammation can reprogram DNA damage responses, with broad implications for immunity and cancer biology.

## Materials and Methods

### Cell lines and generation of clonal population

MCF-7 cells were cultured in RPMI medium (Thermo-Fisher, cat 11835105), while HCT-116 and U2OS cells were cultured in DMEM medium (Thermo-Fisher, cat 31053044); both media were supplemented with 10% of heat-inactivated Fetal Bovine Serum (FBS, Thermo-Fisher, cat 10270106), 100 units/mL penicillin-streptomycin (Thermo-Fisher, cat. 15140122) and 2 mM L-Glutamine (Gibco, Thermo-Fisher, cat. 25030081). Cells were cultured at 37°C in a humidified incubator with 5% CO_2_. For regular maintenance, cells were split every 3-4 days through trypsinization (Thermo-Fisher, cat 11835105), seeded into 75 cm flasks or p100 dishes, and subjected to weekly testing for mycoplasma contamination using a PCR-based assay.

To generate clonal populations, MCF-7 cells were harvested by 1x Trypsin and counted. A final concentration of 1 cell/100 μL was achieved by serial dilutions and 100 μL were pipetted from a reservoir to a 96-well plate. The plate was screened for single colonies in the following weeks, growing colonies were then expanded and screened for p53/p65 expression via Western Blot.

### p53-GFP knock-in

CRISPR/Cas9 gene editing was used to create p53-GFP knock-in (KI) cell lines. p53 KI MCF-7 cells were generated using four plasmids encoding the Cas9-D10A nickase, two guide RNAs (sgRNA1 and sgRNA2), and a repair vector carrying both the GFP gene and a neomycin resistance gene, flanked by two p53 homology arms (∼800 bp). The sequences targeted by sgRNA1 and sgRNA2 on the p53 genomic locus were 5′-GATGACATCACATGAGTGAG-3′ and 5′-CAGCCACCTGAAGTCCAAAA-3′, respectively. MCF-7 cells seeded on a 6-well plate were co-transfected with the four plasmids (625 ng of each vector per well) using Lipofectamine 3000 (Thermo-Fisher, cat. L3000008) and following the manufacturer’s protocol. After transfection, cells were left to recover a few days before two rounds of antibiotic selection (mild, 4 days and long, ∼15 days) with geneticin (G418 Sulfate, Thermo-Fisher, cat. 10131027, used at 800 μg/mL). Cells were next seeded at single cell per well (96 well plate) to isolate single clones. The KI clone used in this study was functionally tested for nuclear localization and abundance of p53-GFP, for the absence of untagged p53, and for the capability to induce the p53 target gene *CDKN1A*.

### Plasmids production and lentiviral infection of p65-mScarlet

Plasmids were produced via bacterial transformation. Each plasmid was diluted to a final concentration of 2-5 ng/mL in H_2_O and the quantity used for each transformation was 15 ng. Each solution was added to the “top 10” bacteria (Thermo-Fisher, cat C404003) and then left in ice for 30 minutes. Bacteria were then subjected to heat-shock (2 minutes, 42°C) and we let them recover in ice for 10 minutes. 1 mL of Lysogeny broth (LB, without antibiotic) was added to bacteria, incubated then for 1 hour, while mildly shaking (300 rpm) at 37°C. Tubes were then centrifuged at 4000 rpm for 5 minutes, the supernatant was discarded and the bacteria pellet was suspended in 400 μL of LB. 200 μL were then spread with a spatula on a plate coated with Ampicillin-resistant LB-agar. Plates were then left upside down overnight at 37°C. The day after, one bacterial colony from the plate was added to a flask containing 120 mL of LB with 1:1000 diluted Ampicillin, and then left on a stirring platform overnight at 37°C. The following day, MIDI-Prep protocol (NucleoBond Xtra EF plasmid purification, Carlo Erba, cat FC140420L) was performed to extract plasmids, that were then quantified via UV spectrophotometry.

The lentivirus for p65-mScarlet was produced in HEK 293T cells. 3×10^6^ HEK 293T cells, cultured in DMEM supplemented with 10% FBS, 1x L-glutamine and 1x pen/strep, were seeded in a p100 dish the day before transfection. The following day, the medium was replaced 2 hours before transfection with 9 mL of DMEM and plasmids, HBS 2X (used volume for p100 plate = 500 μL) and CaCl_2_ (2.5 M, used volume for p100 plate = 50 μL) were thawed before transfection. The protocol relies on three plasmids: p65-mScarlet construct (12000 ng), VSV-G (7000 ng, Addgene, cat 8454) and psPAX2 (11000 ng, Addgene, cat 12260). Each plasmid was added at the working concentration in H_2_O. After 5 minutes at room temperature (RT), CaCl_2_ was added to the master mix. After 5 minutes at RT, HBS is added dropwise, while bubbling the master mix with a 2 mL pipette. The mix was then added dropwise to the plate with HEK 293T cells. The medium was replaced 14 hours after transfection and the supernatant containing the virus was collected, filtered with a 0.22 m syringe filter and added to MCF-7 cells 30 hours after the medium change. Infected cells were incubated at 37°C and 5% CO_2_ and were split 3-4 times with BSL 2 laboratory procedures for at least 2 weeks, after which the virus became inactivated.

### Generation and validation of p65 knock-out MCF-7 p53-GFP cell line

To generate p65 KO MCF-7 cells, a commercial CRISPR/Cas9-based gene editing KO kit was employed (Santa Cruz Biotechnology, cat. sc-400004) including constructs expressing Cas9-D10A nickase, two sgRNAs and a GFP marker transiently expressed for selection. MCF-7 p53-GFP cells were transfected with UltraCruz Transfection Reagent (sc-395739) following the manufacturer’s instructions. The day after transfection, puromycin 1:10000 was added to cells and then cells were sorted for GFP two days after puromycin addition. Single clones were isolated and expanded from the sorted population as described.Successful knock-out of the p65-null clone employed in this work was confirmed by the absence of p65 at the protein level by Western Blot (using these anti-NF-κB rabbit antibodies: monoclonal Cell-Signalling/Euroclone, cat BK8242S, 1:2000 dilution; polyclonal Sigma-Aldrich, cat PC138-100UL, 1:2000 dilution) and the downregulation of a p65 target gene (*NFKBIA*) at the RNA level by qPCR.

### RNA extraction and RT-qPCR

Cells were cultured on 10 cm dishes up to 80-90% of confluency. In time-points post-treatment of interest, cells were lysed in 500 μL of TRIzol reagent (Thermo-Fisher, cat. 15596018) to extract the total RNA. Lysates were then purified using silica membrane columns (NucleoSpin RNA Plus, Machery-Nagel, Carlo Erba, cat FC140955N), thus extracting the RNA that was quantified and tested for purity by NanoDrop fluorometer (Thermo-Fisher). For each sample, 2 μg of RNA underwent reverse transcription to cDNA using the High-Capacity cDNA Reverse Transcription Kit (Thermo-Fisher cat. 4368814). Real-time qPCR analysis was then performed to assess the expression of p53/NF-κB target genes. The reaction solution was prepared as follows: 8 μl of cDNA solution (5 μL of cDNA 1:100 diluted + 3 μL of nuclease-free H2O) and 5 μL of Oligo Solution, composed by 10 μL of SYBR Green mix (Roche, LightCycler 480 I Master) and 2 mL of primer mix (100 μM of forward and reverse primers). GAPDH was used as an internal standard for cDNA normalization.

### Immunofluorescence

MCF-7/U2OS/HCT-116 cells were seeded on coverslips and grown up to 80% confluency. Cells were treated according to the experimental design (γ-irradiation, Nutlin3a, TNF-α, IL-1β or their combination), washed once in PBS and fixed in 4% paraformaldehyde (PFA) for 10 minutes at room temperature. After fixation, cells underwent quenching with 50 mM NH_4_Cl to reduce background, permeabilization with 0.1% Triton X-100 (Sigma Aldrich) and blocking at RT for 45 minutes in PBS supplemented with 20% FBS, 0.05% Tween 20 (Sigma Aldrich, cat. P2287) and 5% BSA (Sigma Aldrich, cat. A7906). Next, cells were incubated for 1 hour at room temperature with primary antibodies (diluted in 2% BSA) to detect p53 (mouse monoclonal antibody, Santa Cruz cat sc-126, 1:200 dilution), NF-κB (rabbit monoclonal antibody, Cell-Signalling/Euroclone, cat BK8242S, 1:100 dilution) or the DNA damage marker γ-H2AX (rabbit polyclonal γ-H2AX antibody, Abcam cat ab11174, 1:200 dilution). After three PBS washes, cells were incubated with secondary antibodies 1:1000 diluted in 2% BSA (anti-mouse IgG Alexa Fluor 488, Thermo-Fisher, cat A11029; anti-rabbit IgG Alexa Fluor 546, Thermo-Fisher, cat A11035; anti-rabbit IgG Alexa Fluor 647, Thermo-Fisher, cat A32733), for 1 hour at RT in the dark. Cells were washed three times in PBS and incubated in the dark with a DNA staining solution (Hoechst 33342, Thermo-Fisher, cat. H3570, 1:5000 dilution in PBS), for 10 min at RT. After two washes with PBS, coverslips were mounted with Vectashield (Vector Laboratories) and sealed.

p53 and NF-κB protein levels/localization were imaged through a Leica TCS SP5 confocal microscope, using a 20x, 0.5 NA objective, open pinhole (Airy = 3) and laser power, gain, offset respectively equal to: 405 nm = 12%, 784, 0; 488 nm = 14%, 784, 0; 561 nm = 35%, 497, 0. γ-H2AX foci were imaged through a widefield microscope (Olympus IX-81), equipped with a sCMOS camera (physical pixel size = 108.3 nm) and a 60x 1.49 N.A. oil-immersion objective (Olympus Life science, Segrate, IT) (UV led exposure: 100 ms; Blue led exposure: 500 ms; Yellow led exposure = 500 ms), collecting z-stacks using a z-step of 0.5 μm.

### Live cell imaging

Cells were seeded and cultured onto μ-Slide 8-Well Glass Bottom supports (Ibidi, cat 80827), in order to reach 70-80% confluency on the day of the live-cell imaging experiment. Before imaging, nuclei cells were stained with Hoechst 33342 (1:10000 in RPMI, 10 minutes incubation at 37°C and 5% CO_2_); the medium was then replaced with 150 μL of fresh medium. Treatments were prepared at double concentration and 150 μL were directly added to the respective well. We used both Leica TCS SP5 and SP8 confocal microscopes, with an incubation system where cells were stably maintained at 37°C in 5% CO_2_. We used a 20x, 0.5 NA objective, open pinhole (Airy = 3) and 400 Hz of acquisition frequency. p53-GFP and p65-mScarlet were imaged with the 488 nm and 546 nm Argon lasers, respectively, in the SP5 microscope and with a White Light Laser in the SP8 microscope. Hoechst 33342 stained nuclei were imaged with the low energy 405 nm UV diode laser (power = 5-10%). Images were acquired as 16-bit, 1024×1024, thus returning a single LIF file containing all TIFF x-y-t stacks. The frequency of acquisition was decided according to the experimental conditions that we were testing and the number of fields of view per condition, but in order to be able to capture the fast NF-κB dynamics on multiple fields and to avoid oversampling, the acquisition frequency was always set between 6-8 minutes.

### Mathematical modeling of p53 dynamics

The model we used, adapted from the one described and used in (31, 37) is described as follows:

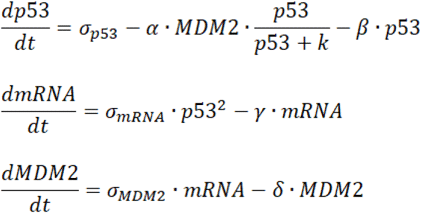

where the variables represent normalized *p53* and *MDM2* protein and *mRNA* concentrations. The parameters σ_p53_, σ_mRNA_ and σ_MDM2_ modulate the rates of p53, MDM2 mRNA and MDM2 synthesis, respectively, α is the parameter modulating the strength of the MDM2-dependent degradation of p53 with hill constant *k*, γ and δ are the degradation rates for mRNA and MDM2 respectively β is a degradation rate of p53 that is not MDM2-mediated. This was added to the model following (38) to produce damped oscillations mimicking the average profiles of p53 activation observed experimentally.

Simulations of p53 dynamics were performed as follows: the system is integrated until it reaches a stable state simulating the “homeostatic values” of the variables, and then parameters were changed to simulate the effect of the treatments. Specifically, following (31), to simulate DNA damage through γ-irradiation, δ was increased by a factor of 3.3 while α was decreased by a factor of 3.0 with respect to the unperturbed value. Instead, to simulate Nutlin3a effect, which blocks MDM2-dependent degradation of p53, α was decreased by three orders of magnitude. In both cases, the remaining parameters are left unchanged. Parameters were chosen after stochastic exploration of parameter space in a way that the simulations following this procedure led to dynamics qualitatively similar to the ones observed experimentally in absence of TNF-α stimulation. For both types of perturbation, to account for the effect of increased *TP53* transcription upon NF-κB-activating stimuli in presence of p53 activating stimuli, we changed the parameters δ and α as above but substituted the p53 synthesis parameter σ_p53_ by a piecewise linear time-dependent function σ_p53_(t), built using the median values of the smRNA-FISH in specific timepoints for the stimuli considered, so its fold change with respect to the unperturbed value matches the experimental fold change of *TP53* mRNA levels. To mimic the effect of NF-κB in absence of p53-activating stimuli, we used σ_p53_(t) extracted from experimental data leaving δ and α unchanged. Numerical integration was performed using MATLAB. Parameters for the model are provided in **Table S1**.

### Transcriptomics and bioinformatic analysis

Total RNA was extracted and assessed for concentration, purity and integrity as described above for qPCR. Sequencing libraries were prepared using the Standard Input mRNA library preparation protocol for NovaSeq 6000, generating 1×100 nt single-end reads, and sequenced on the Illumina NovaSeq 6000 platform (Illumina, San Diego, CA, USA). The quality of raw sequencing reads was assessed using FastQC (v0.12.1; https://www.bioinformatics.babraham.ac.uk/projects/fastqc), and quality control reports were aggregated with MultiQC (v1.28, (50)). Reads were aligned to the human reference genome (GRCh38) using STAR (v2.7.10b, (51)), and gene-level counts were obtained with FeatureCounts (v2.0.8, (52)). The raw count matrix was imported into R (v4.3.2) and analyzed using the DESeq2 pipeline (v1.42.1, (53)). Differentially expressed genes (DEGs) were defined as those with an adjusted p-value < 0.05 and an absolute log2 fold change ≥ 0.58 (corresponding to a fold change > 1.5). Hierarchical clustering was performed using Matlab on gene sets sufficiently expressed (RPKM>1 for all samples) for the gene sets specified, considering genes differentially expressed in at least one condition (applying the Bonferroni-Holmes correction, FDR=0.05). Gene set enrichment analysis where performed providing as inputs sets of t-statistics and gene names and uploading them into https://www.webgestalt.org/ for “Gene set enrichment analysis”, “Homo sapiens”, “pathway”, “KEGG”. Gene sets “γ + TNF wins” and “γ + TNF wins” were defined as sets of genes for which the 16 hours treatments specified led to significantly higher value than for the remaining conditions and the untreated samples. Gene ontology analysis was performed with Enrichr (https://maayanlab.cloud/Enrichr/). Regulatory regions of the genes encoding for p53 were identified as 2000 bp upstream the TSS. For human (*TP53*) and mouse (*trp53*), the UCSC genome browser was using the reference genomes hg38 and mm39, respectively. For zebrafish (*tp53*), the sequence was retrieved from the Ensembl genome browser. Binding sites in the retrieved sequences were explored with JASPAR (https://jaspar.elixir.no/), specifically the presence of the motifs compatible with the matrix profile MA0107.1. In the three species at least 2 binding sites with overlap above 80% were identified.

### smRNA-FISH

Cells were seeded on coverslips and cultured up to 80% of confluency. After treatments, cells were fixed in 4% PFA (10 min at RT) and washed with 135 mM Glycine in PBS for 10 min at RT. smRNA-FISH protocol changed according to the probe. For *CDKN1A* and *NFKBIA* probes (Design Ready Stellaris), cells were incubated with a washing buffer I, composed of 10% saline sodium citrate (SSC) in nuclease-free water (Sigma-Aldrich, cat 95284), for 5 minutes at RT. Then, cells were incubated with a washing buffer II, composed of 10% SSC and 20% deionized formamide in nuclease-free water, for 5 minutes at RT. Next, cells underwent hybridization, by placing coverslips onto a drop (80 μL) of hybridization solution (1 μL of probe in 100 μL of 10% SSC, 10% dextran sulphate and 20% deionized formamide in nuclease-free water) and left overnight in a humidified chamber (37°C). The following day, cells were washed twice with buffer I (30 min per wash, at 37 °C in the dark), once in 10% SCC-20×, and then incubated with a DNA staining solution (Hoechst 33342, 1:10000 diluted in PBS) for 10 min at RT in the dark. For the *TP53* probe (HuluFISH), cells were washed twice with HuluWash buffer (2x SSC, 2M Urea) for 10 minutes each at RT. Cells were then hybridized into a humidified chamber (30°C) by placing the coverslips onto a drop of hybridization solution (0.5 μL of *TP53* probe diluted into 50 μL of 2x SSC, 2M Urea, 10% dextran sulphate, 5x Denhardt’s solution). The following day, cells were washed twice in HuluWash buffer (30 minutes per wash, at 37 °C in the dark) and then incubated with a DNA staining solution (Hoechst 33342, 1:10000 in PBS) for 10 min at RT in the dark. Coverslips were mounted on glass slides with Vectashield (Vector Laboratories) and sealed. 3D z-stack images were collected with a widefield microscope (Olympus IX-81) (z-step = 0.3 μm; UV led exposure: 20 ms; Yellow led exposure (*CDKN1A*): 500 ms; Red led exposure = 500 ms (*NFKBIA*), 100 ms (*TP53*)). Images were then processed with custom-written routines and FISH-Quant (54). Nuclei and cell profiles were segmented via Cellpose and then transformed into FISH-Quant regions-of-interest (ROI) by Python routines. With FISH-Quant, the amount of mature RNA molecules per cell was detected as 3D Gaussian spots exhibiting peak intensity levels above a predetermined threshold, which was kept constant for each analyzed image. The threshold was differently set according to the analyzed probe (*TP53, CDKN1A*, and *NFKBIA*).

### Western Blot

Cells were cultured on p100 dishes up to 80-90% of confluency. According to experimental time-points, dishes were washed once with PBS and lysed in 140 μL RIPA buffer (25 mM Tris HCl pH 7.6, 150 mM NaCl, 1% Sodium deoxycholate, 1% Triton X-100, 2 mM EDTA dihydrate) supplemented with protease inhibitors (Sigma-Aldrich, cat. 4693124001). Samples were next incubated at 4 °C for 15 minutes under constant rotation, centrifuged at 13000 rcf at 4°C and the supernatants were then collected into new vials. Lysates were then quantified via the BCA assay (Thermo-Fisher, cat. 23225). Single lysates were prepared in order to load from 25 to 40 μg of proteins, thus adding H_2_O and 4x Laemmli protein sample buffer (Biorad, cat 1610747, supplemented with 10% of β-mercaptoethanol) up to the final volume. Lysates were then loaded into an 8% SDS-polyacrylamide gels (Lower gel: 4.7 mL H2O, 2.5 mL Resolving Buffer, 6.6 mL Acrylamide, 100 μL ammonium persulfate, 10 μL TEMED; Stacking gel: 2.862 mL H2O, 2.5 mL Upper Buffer, 833 μL Acrylamide, 50 μL ammonium persulfate, 5 μL TEMED); electrophoresis ran at 100 Volts for ∼2-3 hours. Proteins were then transferred to nitrocellulose membranes in cold transfer buffer (25 mM Tris, 192 mM glycine, 20% methanol) through run at 100 Volts for 2 hours (or at 15 V, overnight) at 4°C. Membranes were then exposed to Ponceau solution, washed with TBS-T solution (0.1% Tween20 in TBS: 20 mM Tris base, 137 mM sodium chloride, pH 7.6) and blocked in 5% non-fat dried milk in TBS-T solution for 1 hour at RT on oscillating platform. Membranes were then incubated overnight at 4°C with primary antibodies, all diluted in 5% non-fat dried milk in TBS-T solution (anti-p53 mouse monoclonal antibody, Santa Cruz cat sc-126, 1:3000 dilution; anti-NF-κB rabbit monoclonal antibody, Cell-Signalling/Euroclone, cat BK8242S, 1:2000 dilution; anti-GAPDH rabbit monoclonal Abcam, cat. ab128915, 1:40000 dilution; anti-vinculin mouse monoclonal, Thermo-Fisher, cat. MA5-11690, 1:4000 dilution). Next, membranes were washed three times in TBS-T, 5 min each wash at RT, while shaking, and then incubated for 1 hour at RT with peroxidase-conjugated secondary antibodies (anti-mouse IgG, Cell Signalling, cat. 7076; anti-rabbit IgG, Cell Signalling, cat. 7074) diluted 1:5000 in 5% non-fat dried milk in TBS-T solution. Membranes were then developed using an ECL substrate (Bio Rad, cat. 1705061) and images were acquired with a CCD camera via ChemiDoc MP imaging system. Band intensity was quantified via ImageLab software.

### Cell tracking, Image analysis and time series analysis

Time-courses were processed and analyzed via semi-automatic custom-written ImageJ/FIJI macros and MATLAB scripts. From the global LIF file, single TIFFs, corresponding to the single fields of view, were grouped into separate stacks. Each frame of the stack was then segmented via Cellpose (55), to have masks of both nuclei and entire cell profiles, that were subsequently grouped into new stacks. In MATLAB, each stack was processed one at a time. Each nucleus was associated with its “cytoplasm” according to the level of superimposition between the two masks. Cells detected at the boundary of the image were excluded. p53 and NF-κB mean fluorescence intensity was computed from nuclear masks, while NF-κB N.C.R was computed as follows. The code quantifies the area of both the nucleus and the entire cell (A_nucl_, A_cell_), with respective fluorescence intensities (I_nucl_, I_cell_). Cytoplasm’s fluorescence intensity is computed as:I_cytopl_=((A_cell_*I_cell_-A_nucl_*I_nucl_))/(A_cell_-A_nucl_)). N.C.R. is then calculated as I_nucl_ /I_cytopl_. Each mask was then tracked along the time-course with the Hungarian linker algorithm (56). Single-cell profiles were then stored in several arrays according to the measured entity: p53 nuclear intensity, NF-κB nuclear intensity and N.C.R, nuclei and cytoplasms areas and x-y coordinates in the field of view. Matrices were then concatenated according to the respective treatment and profiles were smoothed with the *smoothdata* MATLAB function (method = Gaussian, window size for p53 = 7 (45), window size for NF-κB = 4). Colorplots were arranged by sorting the profiles in order of increasing maximum value. AUC of p53 and NF-κB profiles (after removing the basal level) were calculated as the integral in the time interval considered, for only those cells that were visible for the entire movie. Peaks of oscillations of p53 dynamics were computed on single-cell profiles reaching at least ⅔ of the length of the acquisition, using the *islocalmax* MATLAB function, thus considering the two peaks with higher prominence. Cells were classified as cells with low and high responses using the top and bottom half AUC NF-κB values, respectively.

To quantify the number of γ-H2AX foci from immunofluorescence samples, we computed the maximum projection of the stacks, and used the ThunderSTORM ImageJ plugin (57) to automatically detect spots of DNA damage (camera setup parameters: offset = 100; photoelectrons per A/D counts = 0.22; basal level = 100; connectivity = 8; threshold for spot detection = 4*std(Wavelet filter) with local maximum method). The algorithm generated a table with the positions of single spots. Cell nuclei were segmented through Cellpose (55). Masks and tables with spots positions were loaded in MATLAB, where a custom-written routine assigned each spot to the respective nucleus and quantified the number of spots per cell. Cells detected at the boundary of the image were excluded.

## Supporting information

Raw Data

## Data and Code Availability

Raw data to reproduce the figures in are provided in Excel format in the zipped file Colombo_etal_2026_RawData.zip. All RNAseq data are available at the NCBI Gene Expression Omnibus (GEO) database, accessible via Series accession number GSE320287. Mathematical modeling can be downloaded at https://github.com/SZambranoS/p53NFkappaB. Bioinformatics analysis tools are available as described in the materials and methods section. Any additional information required to reanalyze the data reported in this paper is available from the corresponding authors upon request.

## AUTHOR CONTRIBUTIONS

Conceptualization: E.C, S.Z and D.M.; Supervision: S.Z. and D.M.; Visualization: E.C., F.G., S.Z and D.M.; Formal Analysis: E.C., F.G., T.H., E.A., S.Z. and D.M.; Investigation: E.C., S.P., A.L., F.G., E.A., T.H., P.F., T.F.,M.M., D.G., A.A., M.E.B., S.Z and D.M.; Writing – Original draft: E.C., M.E.B., S.Z. and D.M.; Writing – Review and editing: E.C., S.P., A.L., F.G., E.A., T.H., P.F., T.F., M.M, D.G., A.A., M.E.B., S.Z and D.M.; Funding acquisition: M.E.B., S.Z. and D.M.

## ACKNOWLEDGEMENTS

We are grateful to the Advanced Light and Electron Microscopy BioImaging Center (ALEMBIC, Ospedale San Raffaele) for the access and the support on the Leica SP8 confocal microscope, and to the Center for Omics Sciences (COSR, Ospedale San Raffaele) for the support on the RNAseq experiments. We thank the members of the Experimental Imaging Center and of the Division of Genetics and Cell Biology (Ospedale San Raffaele) for helpful advice and critical discussions.

## REFERENCES

1. U. Alon, An Introduction to Systems Biology (CRC Press, 2007).

2. S. S. Shen-Orr, R. Milo, S. Mangan, U. Alon, Network motifs in the transcriptional regulation network of Escherichia coli. Nat Genet 31, 64–68 (2002).

3. J. Selimkhanov, et al., Accurate information transmission through dynamic biochemical signaling networks. Science 346, 1370–1373 (2014).

4. F. Bonsignore, S. Pozzi, E. Aloi, D. Mazza, S. Zambrano, Linking signaling dynamics and cell fate decisions through single-cell imaging: evidence and challenges. Front. Cell Dev. Biol. 13 (2025).

5. K. F. Sonnen, et al., Modulation of Phase Shift between Wnt and Notch Signaling Oscillations Controls Mesoderm Segmentation. Cell 172, 1079-1090.e12 (2018).

6. K. Meyer, N. C. Lammers, L. J. Bugaj, H. G. Garcia, O. D. Weiner, Optogenetic control of YAP reveals a dynamic communication code for stem cell fate and proliferation. Nat Commun 14, 6929 (2023).

7. C. Droin, E. R. Paquet, F. Naef, Low-dimensional dynamics of two coupled biological oscillators. Nat. Phys. 15, 1086–1094 (2019).

8. J. Richards, M. L. Gumz, Advances in understanding the peripheral circadian clocks. The FASEB Journal 26, 3602–3613 (2012).

9. G. Carrà, et al., Mechanisms of p53 Functional De-Regulation: Role of the IκB-α/p53 Complex. Int J Mol Sci 17 (2016).

10. P. Ak, A. J. Levine, p53 and NF-κB: different strategies for responding to stress lead to a functional antagonism. FASEB j. 24, 3643–3652 (2010).

11. G. Schneider, O. H. Krämer, NFκB/p53 crosstalk-a promising new therapeutic target. Biochim Biophys Acta 1815, 90–103 (2011).

12. G. Carrà, M. F. Lingua, B. Maffeo, R. Taulli, A. Morotti, P53 vs NF-κB: the role of nuclear factor-kappa B in the regulation of p53 activity and vice versa. Cell. Mol. Life Sci. 77, 4449–4458 (2020).

13. E. Kim, A. Giese, W. Deppert, Wild-type p53 in cancer cells: when a guardian turns into a blackguard. Biochem Pharmacol 77, 11–20 (2009).

14. X. Chen, et al., Mutant p53 in cancer: from molecular mechanism to therapeutic modulation. Cell Death Dis 13, 974 (2022).

15. M. Aqdas, M.-H. Sung, NF-κB dynamics in the language of immune cells. Trends in Immunology 44, 32–43 (2023).

16. M. Karin, F. R. Greten, NF-kappaB: linking inflammation and immunity to cancer development and progression. Nat Rev Immunol 5, 749–759 (2005).

17. F. Colombo, S. Zambrano, A. Agresti, NF-κB, the Importance of Being Dynamic: Role and Insights in Cancer. Biomedicines 6, 45 (2018).

18. M.-K. Choy, et al., High-throughput sequencing identifies STAT3 as the DNA-associated factor for p53-NF-κB-complex-dependent gene expression in human heart failure. Genome Medicine 2, 37 (2010).

19. A. Bisio, et al., Cooperative interactions between p53 and NFκB enhance cell plasticity. Oncotarget 5, 12111–12125 (2014).

20. J. M. Lowe, et al., p53 and NF-κB Coregulate Proinflammatory Gene Responses in Human Macrophages. Cancer Research 74, 2182–2192 (2014).

21. N. Geva-Zatorsky, et al., Oscillations and variability in the p53 system. Molecular Systems Biology 2, E1–E13 (2006).

22. A. Hoffmann, A. Levchencko, M. L. Scott, D. Baltimore, The IkappaB-NF-kappaB signalling module: temporal control and selective gene activation. Science 298, 1241–1245 (2002).

23. A. Jiménez, et al., Time-series transcriptomics and proteomics reveal alternative modes to decode p53 oscillations. Molecular Systems Biology 18, e10588 (2022).

24. S. Zambrano, I. de Toma, A. Piffer, M. E. Bianchi, A. Agresti, NF-kappaB oscillations translate into functionally related patterns of gene expression. eLife 5, e09100–e09100 (2016).

25. R. E. C. Lee, M. A. Qasaimeh, X. Xia, D. Juncker, S. Gaudet, NF-κB signalling and cell fate decisions in response to a short pulse of tumour necrosis factor. Sci Rep 6, 39519 (2016).

26. T. Kull, et al., NfκB signaling dynamics and their target genes differ between mouse blood cell types and induce distinct cell behavior. Blood 140, 99–111 (2022).

27. Q. J. Cheng, et al., NF-κB dynamics determine the stimulus specificity of epigenomic reprogramming in macrophages. Science 372, 1349–1353 (2021).

28. A. Adelaja, et al., Six distinct NFκB signaling codons convey discrete information to distinguish stimuli and enable appropriate macrophage responses. Immunity 54, 916-930.e7 (2021).

29. R. Yang, et al., Cell type–dependent bimodal p53 activation engenders a dynamic mechanism of chemoresistance. Science Advances 4, eaat5077 (2018).

30. A. L. Paek, J. C. Liu, A. Loewer, W. C. Forrester, G. Lahav, Cell-to-Cell Variation in p53 Dynamics Leads to Fractional Killing. Cell 165, 631–642 (2016).

31. M. S. Heltberg, et al., Enhanced DNA repair through droplet formation and p53 oscillations. Cell 185, 4394-4408.e10 (2022).

32. A. Loffreda, et al., Live-cell p53 single-molecule binding is modulated by C-terminal acetylation and correlates with transcriptional activity. Nat Commun 8, 313 (2017).

33. S. Zambrano, et al., First Responders Shape a Prompt and Sharp NF-κB-Mediated Transcriptional Response to TNF-α. iScience 23, 101529 (2020).

34. J. E. Purvis, K. W. K. C. Mock, E. Batchelor, A. Loewer, G. Lahav, p53 Dynamics Control Cell Fate. Science 336, 1440–1444 (2012).

35. H. Wu, G. Lozano, NF-kappa B activation of p53. A potential mechanism for suppressing cell growth in response to stress. Journal of Biological Chemistry 269, 20067–20074 (1994).

36. K. Schumm, S. Rocha, J. Caamano, N. D. Perkins, Regulation of p53 tumour suppressor target gene expression by the p52 NF-κB subunit. EMBO J 25, 4820–4832 (2006).

37. B. Mengel, et al., Modeling oscillatory control in NF-κB, p53 and Wnt signaling. Current Opinion in Genetics & Development 20, 656–664 (2010).

38. R. Lev Bar-Or, et al., Generation of oscillations by the p53-Mdm2 feedback loop: A theoretical and experimental study. Proc. Natl. Acad. Sci. U.S.A. 97, 11250–11255 (2000).

39. E. W. Martin, A. Pacholewska, H. Patel, H. Dashora, M.-H. Sung, Integrative analysis suggests cell type–specific decoding of NF-κB dynamics. Science Signaling 13, eaax7195 (2020).

40. H. Wajant, K. Pfizenmaier, P. Scheurich, Tumor necrosis factor signaling. Cell Death Differ 10, 45–65 (2003).

41. M. Fischer, Census and evaluation of p53 target genes. Oncogene 36, 3943–3956 (2017).

42. J. Stewart-Ornstein, et al., p53 dynamics vary between tissues and are linked with radiation sensitivity. Nat Commun 12, 898 (2021).

43. Y. Ben-Neriah, M. Karin, Inflammation meets cancer, with NF-κB as the matchmaker. Nat Immunol 12, 715–723 (2011).

44. X. Chen, et al., DNA damage strength modulates a bimodal switch of p53 dynamics for cell-fate control. BMC Biol 11, 73 (2013).

45. Y.-H. Wang, et al., Rapid recruitment of p53 to DNA damage sites directs DNA repair choice and integrity. Proc Natl Acad Sci U S A 119, e2113233119 (2022).

46. F. Konrath, A. Mittermeier, E. Cristiano, J. Wolf, A. Loewer, A systematic approach to decipher crosstalk in the p53 signaling pathway using single cell dynamics. PLOS Computational Biology 16, e1007901 (2020).

47. D. Chowdhury, J. Lieberman, Death by a Thousand Cuts: Granzyme Pathways of Programmed Cell Death. Annu Rev Immunol 26, 389–420 (2008).

48. K. De Bosscher, et al., Glucocorticoids repress NF-κB-driven genes by disturbing the interaction of p65 with the basal transcription machinery, irrespective of coactivator levels in the cell. Proceedings of the National Academy of Sciences 97, 3919–3924 (2000).

49. A. M. Cook, A. M. McDonnell, R. A. Lake, A. K. Nowak, Dexamethasone co-medication in cancer patients undergoing chemotherapy causes substantial immunomodulatory effects with implications for chemo-immunotherapy strategies. OncoImmunology 5, e1066062 (2016).

50. P. Ewels, M. Magnusson, S. Lundin, M. Käller, MultiQC: summarize analysis results for multiple tools and samples in a single report. Bioinformatics 32, 3047–3048 (2016).

51. A. Dobin, et al., STAR: ultrafast universal RNA-seq aligner. Bioinformatics 29, 15–21 (2013).

52. Y. Liao, G. K. Smyth, W. Shi, featureCounts: an efficient general purpose program for assigning sequence reads to genomic features. Bioinformatics 30, 923–930 (2014).

53. M. I. Love, W. Huber, S. Anders, Moderated estimation of fold change and dispersion for RNA-seq data with DESeq2. Genome Biol 15, 550 (2014).

54. F. Mueller, et al., FISH-quant: automatic counting of transcripts in 3D FISH images. Nat Methods 10, 277–278 (2013).

55. C. Stringer, T. Wang, M. Michaelos, M. Pachitariu, Cellpose: a generalist algorithm for cellular segmentation. Nat Methods 18, 100–106 (2021).

56. G. Careccia, et al., Exploiting Live Imaging to Track Nuclei During Myoblast Differentiation and Fusion. Journal of Visualized Experiments (JoVE) e58888 (2019). 10.3791/58888.

57. M. Ovesný, P. Křížek, J. Borkovec, Z. Švindrych, G. M. Hagen, ThunderSTORM: a comprehensive ImageJ plug-in for PALM and STORM data analysis and super-resolution imaging. Bioinformatics 30, 2389–2390 (2014).

