## Supplementary Information for "NF-κB transcriptionally enhances p53 accumulation dynamics hampering DNA repair"

**This document includes:**

- 4 Supplementary Figures;
- 1 Supplementary Table;
- 2 Supplementary Movies captions.

#### SUPPLEMENTARY FIGURES

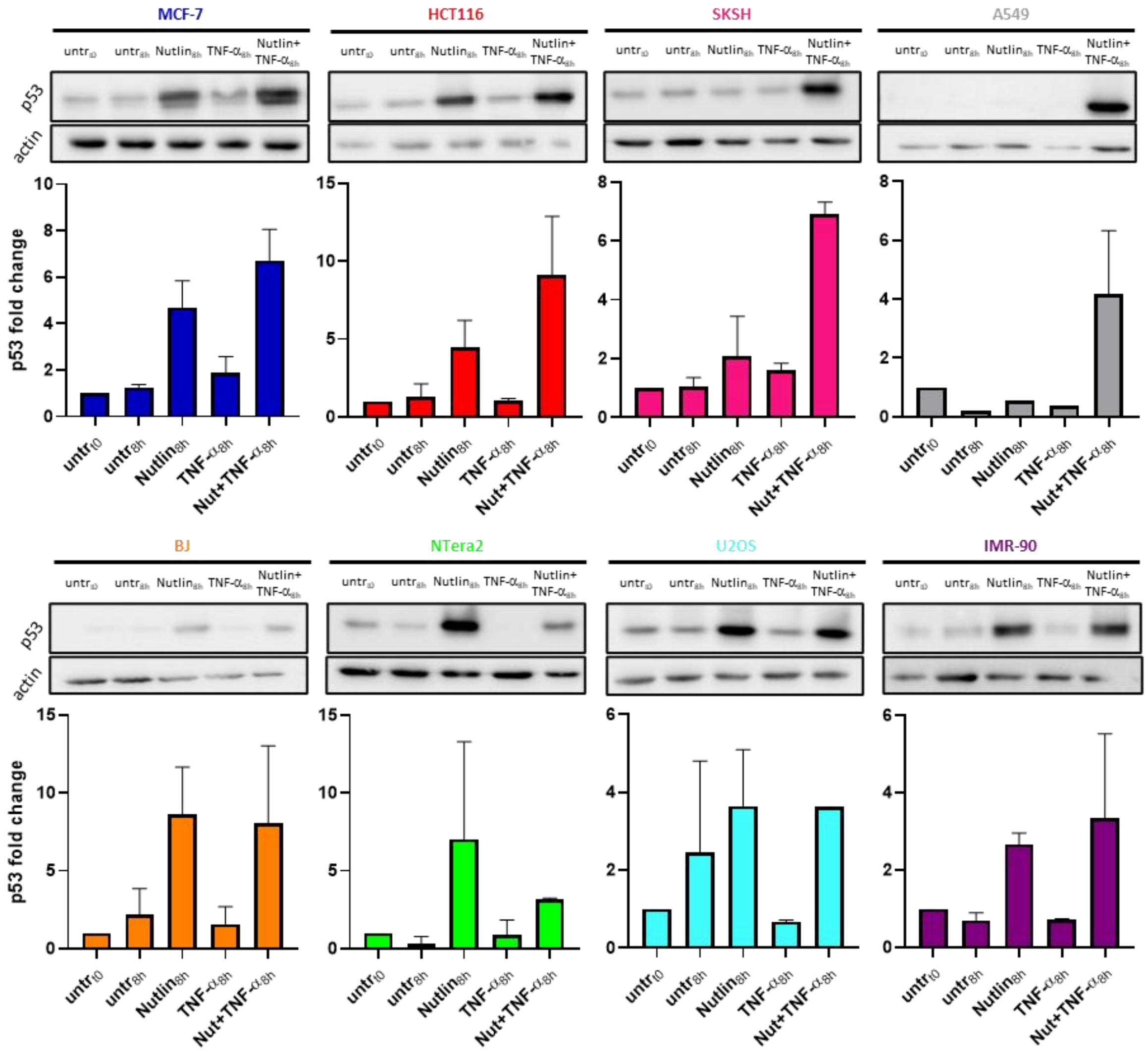


**Figure S1. Co-treatment with TNF-α lead to enhanced p53 response upon Nutlin3a for different cell lines**. Western-blot against p53 and quantification for cells upon Nutlin3a, TNF-α and Nutlin3a+TNF-α treatments for untreated cells and 8 hour stimulated cells of MCF-7, HCT-116, SKSH, A549, BJ, NTera, U2OS and IMR-90 cell lines (n = 2 replicates for each cell line).

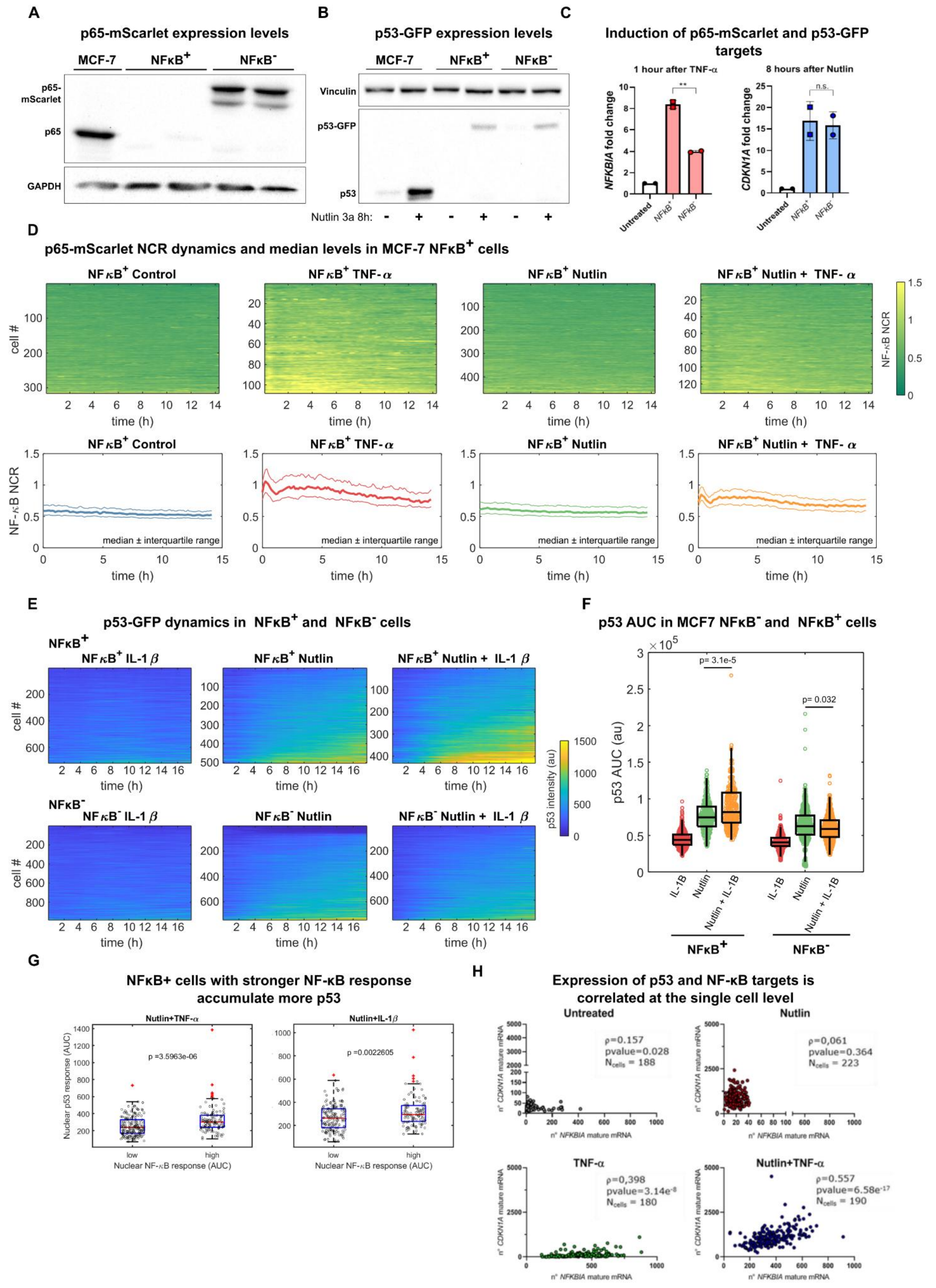


**Figure S2 (Legend in next page)**

**Figure S2. Experimental data supporting NF-κB dependent enhancement of p53 response upon Nutlin3a when co-treated with cytokines.** **A** Western-blot showing the effective knock-out of p65 in NFκB⁻ cells and expression of mScarlet-p65 in NFκB⁺ cells. **B** Western-blot showing activation of p53 upon Nutlin3a for NFκB⁻ and NFκB⁺ cells. **C** *NFKBIA* and *CDKN1A* expression by RT-qPCR upon TNF-α or Nutlin3a in NFκB⁺ and NFκB⁻ cells. **D** Colorplots of NF-κB activation dynamics for hundreds of NFκB⁺ single cells upon different treatments and representation of the median, first and third quartile upon Nutlin3a, TNF-α and Nutlin3a+TNF-α. **E** Colorplots of p53 activation dynamics hundreds of NFκB⁺ and NFκB⁻ single cells upon IL-1ß, Nutlin3a, Nutlin3a+IL-1ß treatments and representation of the median, first and third quartile. **F** Quantification and comparison of the p53 AUC of NFκB⁺ and NFκB⁻ cells upon IL-1ß, Nutlin3a and Nutlin3a+IL-1ß (n = 720, 504, 432 respectively for NFκB⁺ and n=987, 744, 759 cells respectively for NFκB⁻ cells. Statistical test: Kruskal-Wallis, only relevant comparison shown for clarity). **G** p53 response of cells classified as high and low NF-κB response upon Nutlin3a+TNF-α and Nutlin3a+IL-1ß treatments (Statistical test: Kolmogorov-Smirnov). **H** expression of *NFKBIA* and *CDKN1A* quantified via smRNA-FISH for untreated, Nutlin3a, TNF-α and Nutlin3a+TNF-α 8 hour treatments (each dot represents a single cell, statistical test: Pearson correlation).

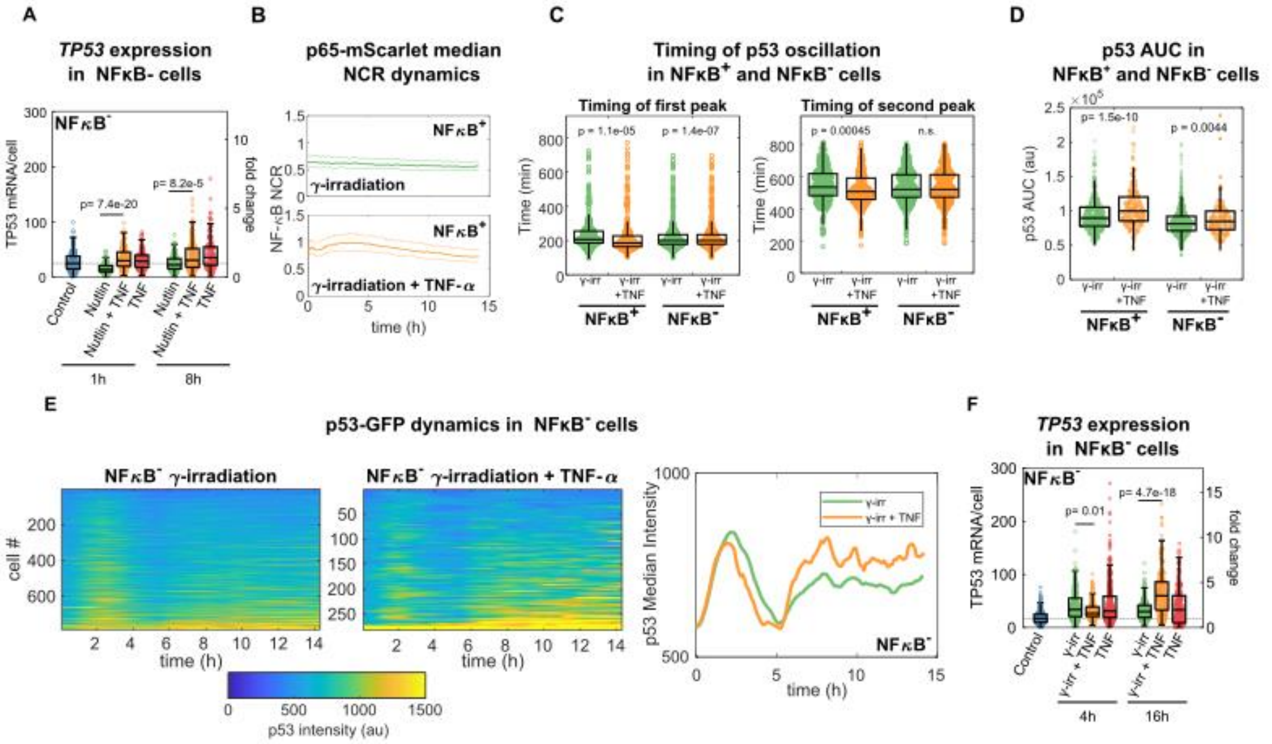


**Figure S3. The perturbation of the p53 oscillatory response upon cytokine stimulation is p65-dependent.** **A** *TP53* expression determined by smRNA-FISH upon Nutlin3a and Nutlin3a+TNF-α treatments and quantification in NFκB⁻ cells (n = 250, 167, 160, 155, 152, 173, 142 cells for control, Nutlin3a 1h, Nutlin3a+TNF-α 1h, TNF-α 1h, Nutlin3a 8h, Nutlin3a+TNF-α 8h, TNF-α 8h, respectively. Statistical test: Kruskal-Wallis, only relevant comparison shown for clarity). **B** Median, first and third quartile for NF-κB activation dynamics upon γ-irradiation, TNF-α and γ-irradiation+TNF-α. **C** Oscillatory features of p53 dynamics for NFκB⁺ and NFκB⁻ cells upon γ-irradiation and γ-irradiation +TNF-α. **D** AUC of p53 dynamics for NFκB⁺ and NFκB⁻ cells upon γ-irradiation and γ-irradiation +TNF-α (statistical test Kruskal-Wallis, Bonferroni-corrected for multiple testing, only relevant comparisons shown for clarity). **E** Colorplots representing p53 activation of hundreds of NFκB⁻ single cells upon γ-irradiation and γ-irradiation +TNF-α and median activation of p53 in NFκB⁻ cells upon γ-irradiation and γ-irradiation +TNF-α. **F** Quantification of *TP53* expression assayed via smRNA-FISH in NFκB⁻ cells upon TNF-α, γ-irradiation and γ-irradiation +TNF-α (n = 176, 131, 184, 282, 201, 153, 146 cells for control, γ-irradiation 4h, γ-irradiation+TNF-α 4h, TNF-α 4h, γ-irradiation 16h, γ-irradiation+TNF-α 16h, TNF-α 16h respectively, statistical test: Kruskal-Wallis, only relevant comparisons shown for clarity).

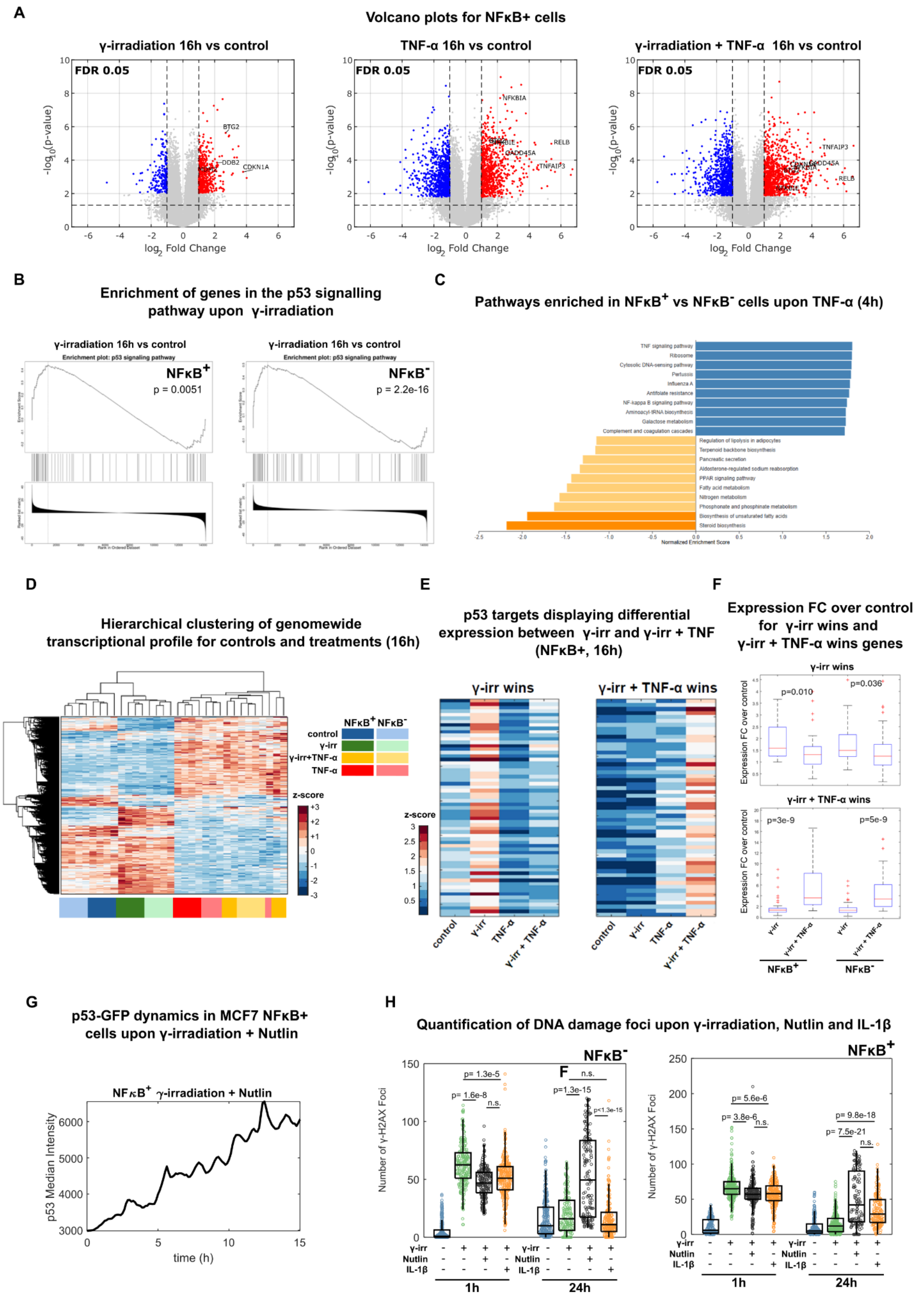


Figure S4 (Legend in next page)

**Figure S4. Characterization of the transcriptional effect of combined treatments and perturbation of DNA repair upon IL-1ß.** **A** Vulcano plots showing activation of different treatments against untreated conditions for NFκB⁺ cells, representative genes are shown. **B** Enrichment of the p53 pathway genes upon γ-irradiation for NFκB⁺ and NFκB⁻ cells. **C** Enriched pathways in NFκB⁺ samples treated with 4h TNF-α versus NFκB⁻ samples under the same treatment. **D** Hierarchical clustering for differentially expressed genes between untreated samples and samples for each treatment at 16h for NFκB⁺ and NFκB⁻ cells. **E** Identification of the "γ + TNF wins" and "γ wins" genes for NFκB⁺ cells. **F** Fold change expression after 16h of the genes of the "γ wins" and "γ + TNF wins" genes sets upon γ-irradiation and γ-irradiation +TNF-α for NFκB⁺ and NFκB⁻ cells. **G** Median non-oscillatory p53 dynamics resulting from the co-treatment γ-irradiation +Nutlin3a. **H** Quantification of the number of gamma-H2AX foci for γ-irradiation 1h, γ-irradiation+Nutlin3a 1h, γ-irradiation+IL-1ß 1h, control 24h, γ-irradiation 24h, γ-irradiation+Nutlin3a 24h, γ-irradiation+IL-1ß, 24h for NFκB⁺ cells (n = 180, 270, 213, 232, 211, 267, 115, 144 cells, respectively, statistical test: Kruskal-Wallis, only relevant comparisons shown for clarity) and NFκB⁻ cells (n = 329, 231, 213, 231,184 , 156, 148, 200 cells, respectively, statistical test: Kruskal-Wallis, only relevant comparisons shown for clarity).

### SUPPLEMENTARY TABLE

| Parameter | Value (“homeostatic values”) |
|---|---|
| $\sigma_{p53}$ | 0.01 $min^{-1}$ |
| $\alpha$ | 0.048 $min^{-1}$ |
| $k$ | 0.1 |
| $\sigma_{mRNA}$ | 0.016 $min^{-1}$ |
| $\gamma$ | 0.016 $min^{-1}$ |
| $\sigma_{MDM2}$ | 0.032 $min^{-1}$ |
| $\delta$ | 0.0048 $min^{-1}$ |
| $\beta$ | 0.005 $min^{-1}$ |

**Table S1.** Parameters used for the mathematical modelling of p53 dynamics.

#### SUPPLEMENTARY MOVIES CAPTIONS

**Supplementary Movie S1.** Exemplary live-cell imaging NFκB$^{+}$ MCF-7 cells, upon Nutlin3a (left) and Nutlin3a+TNF-α (right). p65-mScarlet is shown in red, p53-GFP is shown in green, Hoechst 33342 is shown in blue.

**Supplementary Movie S2.** Exemplary live-cell imaging NFκB$^{+}$ MCF-7 cells, upon γ-irradiation (left) and γ-irradiation +TNF-α (right). p53-GFP is shown in green, Hoechst 33342 is shown in blue.
